# Nuclear receptor interaction protein (NRIP) as a novel actin-binding protein involved in invadosome formation for myoblast fusion

**DOI:** 10.1101/2022.06.14.496213

**Authors:** Hsin-Hsiung Chen, Ya-Ju Han, Tung-Chien Wu, Won-Shin Yen, Tzu-Yun Lai, Po-Han Wei, Li-Kai Tsai, Hsing-Jung Lai, Yeou-Ping Tsao, Show-Li Chen

## Abstract

To investigate the role of nuclear receptor interaction protein (NRIP) in myoblast fusion, both the primary myoblasts from muscle-specific NRIP-knockout mice and NRIP-null C2C12 cells (KO19 cells) exhibited a significant deficit in the fusion index during myogenesis; on the other hand, overexpressed NRIP in KO19 cells could rescue myotube formation. Furthermore, NRIP was found to interact with actin directly and reciprocally that is an invadosome component for myoblast fusion. Endogenous NRIP colocalized with components of invadosome such as F-actin, Tks5, and cortactin at the tips of cells during C2C12 differentiation, and exogenous NRIP was enriched with actin at the tip of attacking cells during myogenic fusion, implying that NRIP is a novel invadosome component. Using time-lapse microscopy and cell–cell fusion assays further confirmed NRIP directly participates in cell fusion through actin. Moreover, to map the domain of NRIP–actin binding, NRIP interacted with actin either through WD40 domains directly for binding or indirectly through the IQ domain for α-actinin 2 binding with actin. NRIP with actin binding was strongly correlated with invadosome formation and myotube fusion. Collectively, NRIP acts as a novel actin-binding protein through its WD40 or the IQ to form invadosomes that trigger myoblast fusion.

## Introduction

The nuclear receptor interaction protein (NRIP) (also named DCAF6 and IQWD1) consists of 860 amino acids, with seven WD40 repetitions and one IQ motif (Chang et al., 2011; Tsai et al., 2005). NRIP is a multifunctional protein interacting with various proteins to execute its biological function in different cells or tissues. For example, NRIP is an androgen receptor (AR)-binding protein mediating AR-targeted gene expression (Chen et al., 2008). NRIP, like DDB2, belongs to the DCAFs family [DDB1-Cul4 associated factors, (Jin et al., 2006)], which can interact with CUL4 and DDB1 to prevent DDB2 from binding with AR to proceed with the DDB2-mediated ubiquitination and degradation of AR through the CUL4-DDB1 E3 ligase complex, causing prostate cancer growth (Chen et al., 2017). NRIP can bind with calmodulin (CaM) through its IQ motif to activate downstream calcineurin (CaN) and calmodulin kinase II (CaMKII) signaling (Chang et al., 2011), thereby regulating Ca^2+^ homeostasis by releasing Ca^2+^ to cytoplasm and uptaking internal Ca^2+^ to the sarcoplasmic reticulum (SR) to regulate muscle contraction (Chen et al., 2015).

NRIP global knockout mice (NRIP KO) exhibit muscle dysfunction but remain vital (Chen et al., 2015). Furthermore, muscle-restricted NRIP-knockout (cKO) mice also show muscle dysfunction and retrograde motor neuron degeneration with an abnormal architecture of their neuromuscular junction (NMJ) (Chen et al., 2018). Additionally, NRIP binding to α-actinin 2 (ACTN2) at the Z-disc can facilitate ACTN2-mediated F-actin bundling; hence, the deficiency of NRIP causes the disruption of sarcomere structural integrity, leading to cardiomyopathy (Yang et al., 2019). Of note, NRIP can be induced after muscle injury, and a deficiency of NRIP delays the repair of myogenesis (Chen et al., 2015). Hence, NRIP is essential for adult muscle regeneration.

Myogenesis is the process of developing skeletal muscle during embryonic development and regeneration after injury. In the beginning, myogenic progenitor cells are activated from quiescence to become myoblasts, followed by the proliferation and differentiation of myoblasts into myocytes, myocyte fusion, and myotube formation. Satellite cells located between the basal lamina and the sarcolemma of muscle fibers are among the postnatal myogenic stem cells responsible for the growth, repair, and maintenance of skeletal muscles (Chen & Goldhamer, 2003; Zammit, Partridge, et al., 2006). In mature skeletal muscles, most satellites are quiescent. When responding to injury, quiescent stem cells are activated to initiate a regeneration program (De Bari et al., 2003). The injured muscles release the hepatocyte growth factor (HGF), the serum response factor (SRF), and the fibroblast growth factor 2 (FGF2) to recruit and activate quiescent paired box transcription factor 7 (Pax7)-expressing satellite cells toward lesion sites (Aline & Sotiropoulos, 2012; Miller et al., 2000; Yablonka-Reuveni et al., 1999; Yin et al., 2013). The activated satellite cells proliferate and differentiate into myoblasts that co-express Pax7 and myogenic differentiation factors (such as MyoD and Myf5) (Bentzinger et al., 2012; Zammit, Relaix, et al., 2006). The committed myoblasts expressing only myogenic differentiation factor (MyoD) but not Pax7 then differentiate into myogenin (MyoG)-expressing myocytes that fuse with each other or with the existing damaged fibers to form multinucleated myofibers for muscle repair and replacement (Ishibashi et al., 2005; Olguin et al., 2007; Yin et al., 2013). As myotubes mature, they undergo secondary fusion to form myofibers that are bundled to take the shape of skeletal muscles (Bentzinger et al., 2012; Berkes & Tapscott, 2005; Kang & Krauss, 2010; Knight & Kothary, 2011).

Myoblast fusion plays a fundamental role in forming a nucleated syncytium to execute muscle function. Actin polymerization/depolymerization regulates the fusion process between fusion-competent myoblasts and founder cells during myogenesis. The mechanisms of muscle cell fusion consist of highly ordered steps. First, cell adhesion molecules responsible for muscle cell adhesion, such as the immunoglobulin (Ig) superfamily, integrins, and cadherins; then, muscle cells extend actin-based lamellipodium and filopodium to recognize, adhere to, and contact neighboring muscle cells. Second, two fusion partners rearrange the actin cytoskeleton to form an F-actin-enriched, invasive podosome-like structure (PLS, also called an invadosome) for invasion (Abmayr & Pavlath, 2012; Kim et al., 2015). Several actin-binding proteins functioning in actin cytoskeleton remodeling are important for myoblast fusion as they facilitate the formation of actin-based protrusion and by assist in actin polymerization to provide a positional cue for the transport of prefusion vesicles (Kim et al., 2007; Nowak et al., 2009; Richardson et al., 2007). For example, the actin-related protein 2/3 (Arp2/3) complex is essential for actin polymerization and promotes the formation of new actin filaments by branching into preexisting actin filaments. The suppressor of cAMP activator *Drosophila* (SCAR) is a pentameric actin-remodeling complex that promotes actin polymerization via the activation of the Arp2/3 complex (Berger et al., 2008).

NRIP can be induced after muscle injury, and NRIP-KO mice reveal more small-sized myofibers compared to wild-type (WT) mice after muscle injury, which indicates that NRIP is essential for myogenesis to execute muscle regeneration (Chen et al., 2015). NRIP can directly interact with ACTN2, which is bundled with actin and required for myoblast fusion (Yang et al., 2019). In this study, we investigated NRIP’s functional role in myogenesis differentiation and myoblast fusion. Our results showed that NRIP affected myogenesis differentiation and also acted as a novel actin-binding protein, forming an invadosome-like structure for myoblast fusion, which resulted in large-sized myofibers.

## Results

### NRIP expression profile during muscle development

Previously, the NRIP transcript was found to be ubiquitously expressed in normal adult human tissues and highly expressed in skeletal muscle and testis (Tsai et al., 2005). In adult mice, the NRIP is also highly expressed in muscle, brain, and testis tissues (Chen et al., 2015; Chen et al., 2018). NRIP functions in muscle contraction (Chen et al., 2015). To further examine the NRIP expression profile in muscle development, NRIP RNA *in situ* hybridization was performed with two DIG-labeled antisense NRIP RNA probes and a sense probe. NRIP RNA was highly expressed in the limb bud, heart, somites, and forebrain of mouse embryos (Figure 1A). NRIP can be detected in the mouse limb bud, heart, and brain at E14.5, and NRIP level was high in the limb bud but was not significantly different from the levels at the brain and heart (Figure 1B). To specifically demonstrate muscle NRIP expression during muscle development, the NRIP level peaked at postnatal day 1 and was maintained at that level at age 6 weeks (Figure 1C). In sum, muscle NRIP expression was higher in the postnatal than in the embryonic period.

**Figure 1.**
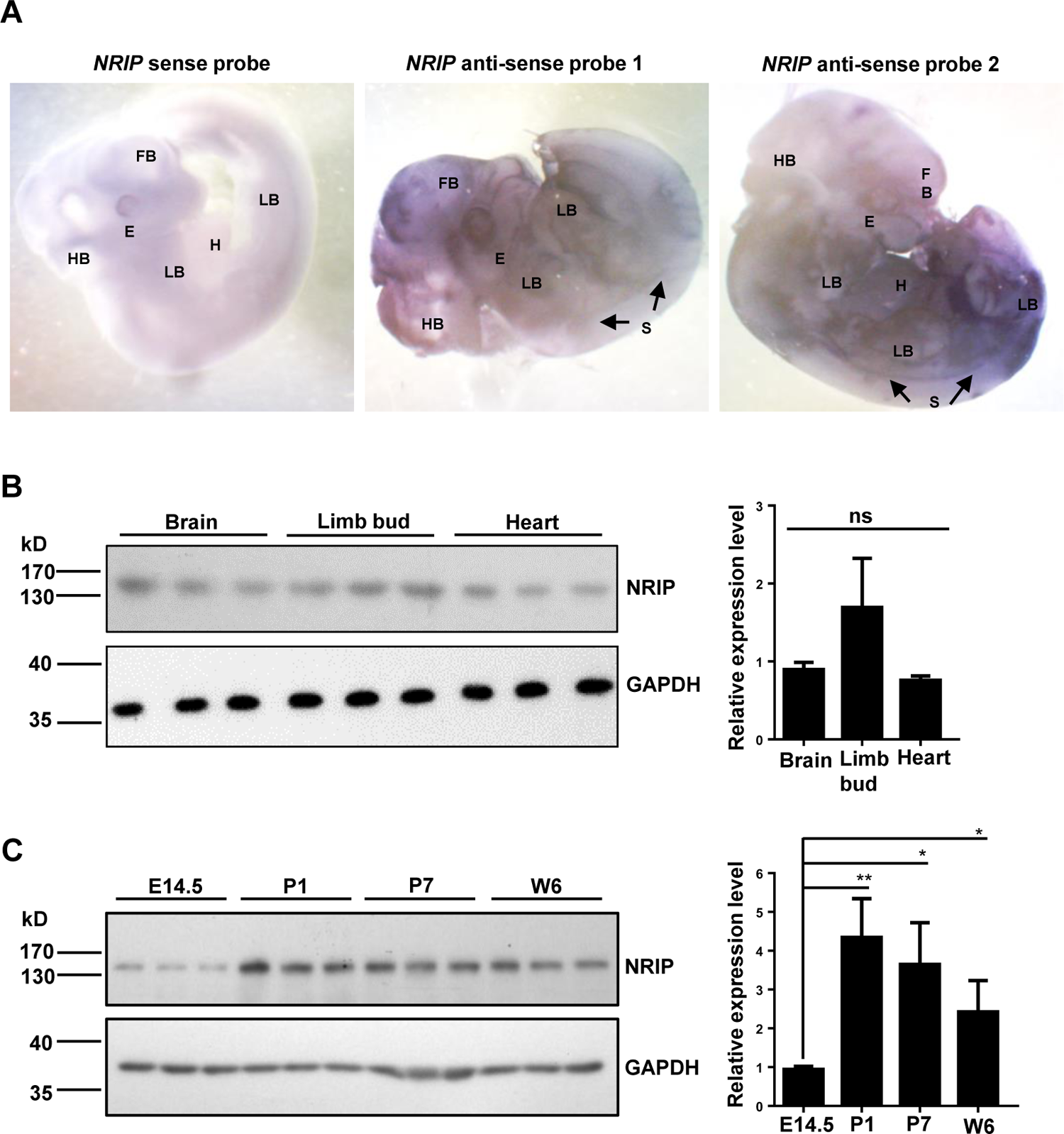
NRIP expression profile in muscle development. (A) NRIP RNA expression in mouse embryos by RNA *in situ* hybridization. E11.5 mouse embryos were hybridized with two different digoxigenin (DIG)-labeled NRIP RNA antisense probes (antisense probe 1-targeted exon 1 to 7, length 696 bp; antisense probe 2-targeted exon 14 to 19; length 578 bp) or with a sense probe (sense probe-targeted exon 1 to 7, length 696 bp) as a negative control, then detected by alkaline phosphatase (AP)-conjugated anti-digoxigenin antibody followed by reaction of purple AP substrate for color development. The dark purple represents positive signals of NRIP transcript expression. FB (forebrain); E (eye); HB (hindbrain); (H) heart; (LB) limb buds; (S) somites (marked by arrow). (B) NRIP was expressed in mouse embryos. Western blot analysis of NRIP expression in brain, limb bud and heart from E14.5 mouse embryos. GAPDH was the loading control. Right: quantification (N=3). (C) The NRIP was significantly expressed in skeletal muscle from newborn mouse. Western blot analysis of muscle NRIP level at embryo and postnatal stages. Skeletal muscles at E14.5, postnatal day 1 (P1), day 7 (P7), and age 6 weeks (W6). Right: quantification (N=3). Data are mean ± SEM by student *t* test. Data are representative of three independent experiments. * \**P* < 0.05, \*\**P* < 0.01 and ns, not significance. See also Figure 1—source data 1 and 2. Source data 1: Numerical data for Figure 1B and 1C. Source data 2: Source WB blot for Figure 1B and 1C.

### The loss of NRIP in primary myoblasts reduces muscle differentiation and fusion

Previously, NRIP KO mice showed small myofibers as compared with WT mice after muscle injury (Chen et al., 2015), which indicates that NRIP is required for adult myogenesis. To examine NRIP’s role in myogenesis, we isolated myoblasts from the hindlimbs of muscle-specific NRIP cKO and WT mice; the isolated myoblasts were then differentiated for 3 days and stained with myosin heavy chain (MyHC) (Shahini et al., 2018) (Figure 2A). The proportion of MyHC^+^ cells with 1 or 2 nuclei was higher in NRIP cKO than WT myoblasts (37.62% vs. 20.15%, *P* < 0.05; 19.44% vs. 9.91%, *P* < 0.01), but that of MyHC^+^ cells containing ≥ 3 nuclei was lower (43.20% vs. 69.83%, *P* < 0.01, Figure 2B), so NRIP is essential for myogenesis.

**Figure 2.**
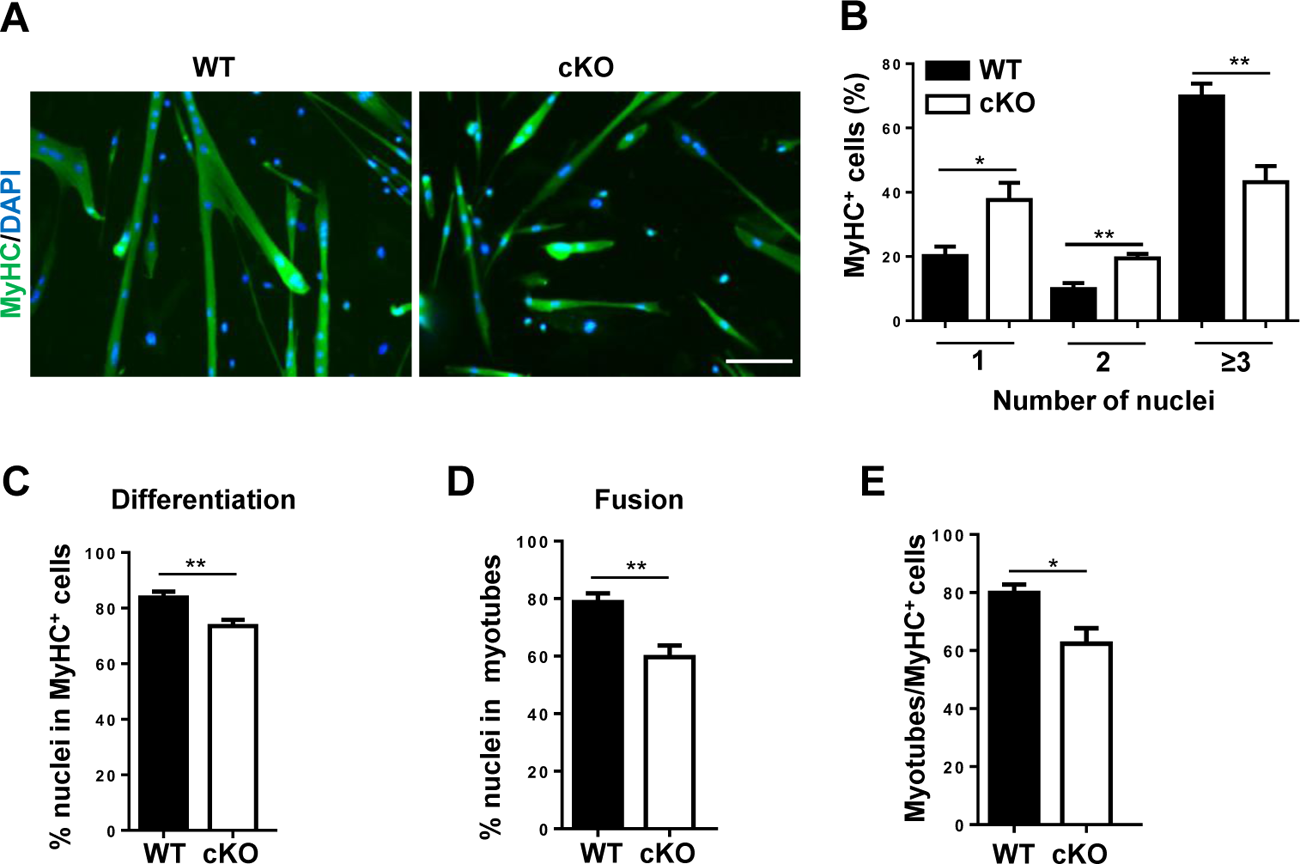
Defective differentiation and fusion indexes of primary myoblasts isolated from NRIP conditional knockout (cKO) mice. (A) Loss of NRIP reduced myotube formation in mouse primary myoblast. Primary myoblasts were dissected from hindlimb muscles from 6-week-old NRIP cKO and wild-type (WT) mice and were seeded on Matrigel-coated dishes. Cells were differentiated for 3 days and stained with anti-myosin heavy chain (MyHC) antibody (green) and DAPI (blue) for nucleus. (B) The distribution of MyHC^+^ cells containing 1, 2 and ≥ 3 nuclei (N=9). (C) Differentiation index calculated as the percentage of nuclei in MyHC^+^ cells (including mononuclear and multinuclear cells) to total nuclei per field (N=9). (D) Fusion index measured by the percentage of nuclei contained in MyHC^+^ myotubes (≥ 2 nuclei) to total nuclei per field (N=9). (E) Percentage of myotubes (≥ 2 nuclei of MyHC^+^ cells) to total MyHC^+^ (N=9). Data are mean ± SEM by student *t* test from 3 separate experiments in which at least three fields of each experiment were randomly measured. \**P* < 0.05, and \*\**P* < 0.01. See also Figure 2— source data 1. Source data 1: Numerical data for Figure 2B to 2E.

Myogenesis consists of two key steps: myoblasts differentiating to myocytes and myocytes fusing together to form a multinucleated myotube (Bentzinger et al., 2012; Berkes & Tapscott, 2005; Kang & Krauss, 2010; Knight & Kothary, 2011). The differentiation index (calculated as the percentage of nuclei in MyHC^+^ cells to total nuclei per field) was lower in NRIP cKO than in WT myoblasts (73.57% vs. 83.78%, *P* < 0.01, Figure 2C). The fusion index, measured as the proportion of nuclei in myotubes (MyHC^+^ cells with ≥ 2 nuclei) to total nuclei, was lower in NRIP cKO myoblasts when compared to WT myoblasts (59.69% vs. 78.78%, *P* < 0.01, Figure 2D). Thus, the NRIP has dual functions for muscle differentiation and myoblast fusion. To demonstrate the NRIP-independent role in myocyte fusion, myotube formation derived from differentiation promotion was excluded; hence, the proportion of myotubes (MyHC^+^ cells with ≥ 2 nuclei) to total MyHC^+^ cells was counted; and showed lower in NRIP cKO than WT mice (62.38% vs. 79.86%, *P* < 0.05, Figure 2E). Thus, NRIP plays an independent role in myoblast fusion. In summary, NRIP is essential for myogenesis differentiation and myocyte fusion.

### NRIP-null C2C12 cells lose myoblast differentiation and fusion

To further investigate the role of NRIP in myogenesis, NRIP-null C2C12 cells were generated by the CRISPR-Cas9 system (Figure 3-figure supplement 1). Targeting NRIP exon 1 by pairs of sgRNAs resulted in frameshift mutation to disrupt NRIP expression (Figure 3B). NRIP-null C2C12 myoblasts were then generated (Figure 3-figure supplement 1C), and KO19 cells were chosen for the following study. We examined the MyHC-positive myocytes by immunofluorescence stain (Figure 3A) and found much less MyHC^+^ cells in KO19 than in C2C12 cells at both 5 and 8 days after differentiation. KO19 cells hardly formed myotubes 5 days after differentiation, but they produced a limited number of myotubes 8 days after differentiation. We measured the differentiation and fusion indexes for KO19 myoblasts at 8 days. The proportion of MyHC^+^ cells containing 1–3 nuclei was higher in KO19 than in C2C12 cells (95.56% vs. 57.92%, *P* < 0.001); cells containing 4–10 nuclei were fewer (4.44% vs. 33.53%, *P* < 0.001), and cells with more than 10 nuclei were absent in KO19 cells in comparison to C2C12 cells (8.55%) (Figure 3C), which indicates impaired multinucleated myotube formation in KO19 myoblasts. Furthermore, when compared to C2C12 cells, KO19 cells showed a reduced differentiation index (6.39% vs. 35.38%, *P* < 0.01, Figure 3D) and fusion index (3.16% vs. 33.67%, *P* < 0.005, Figure 3E). To further demonstrate the NRIP’s independent role in myocyte fusion, the proportion of myotubes (MyHC^+^ cells with ≥ 2 nuclei) to total MyHC^+^ cells was lower in KO19 than WT myoblasts (27.81% vs. 78.70%, *P* < 0.001, Figure 3F), which indicates that NRIP alone can affect myoblast fusion. These results were consistent with the reduced differentiation and fusion indexes for primary myoblasts obtained from NRIP cKO mice (Figure 2). This further confirms that NRIP is responsible for myoblast differentiation and myotube formation.

**Figure 3.**
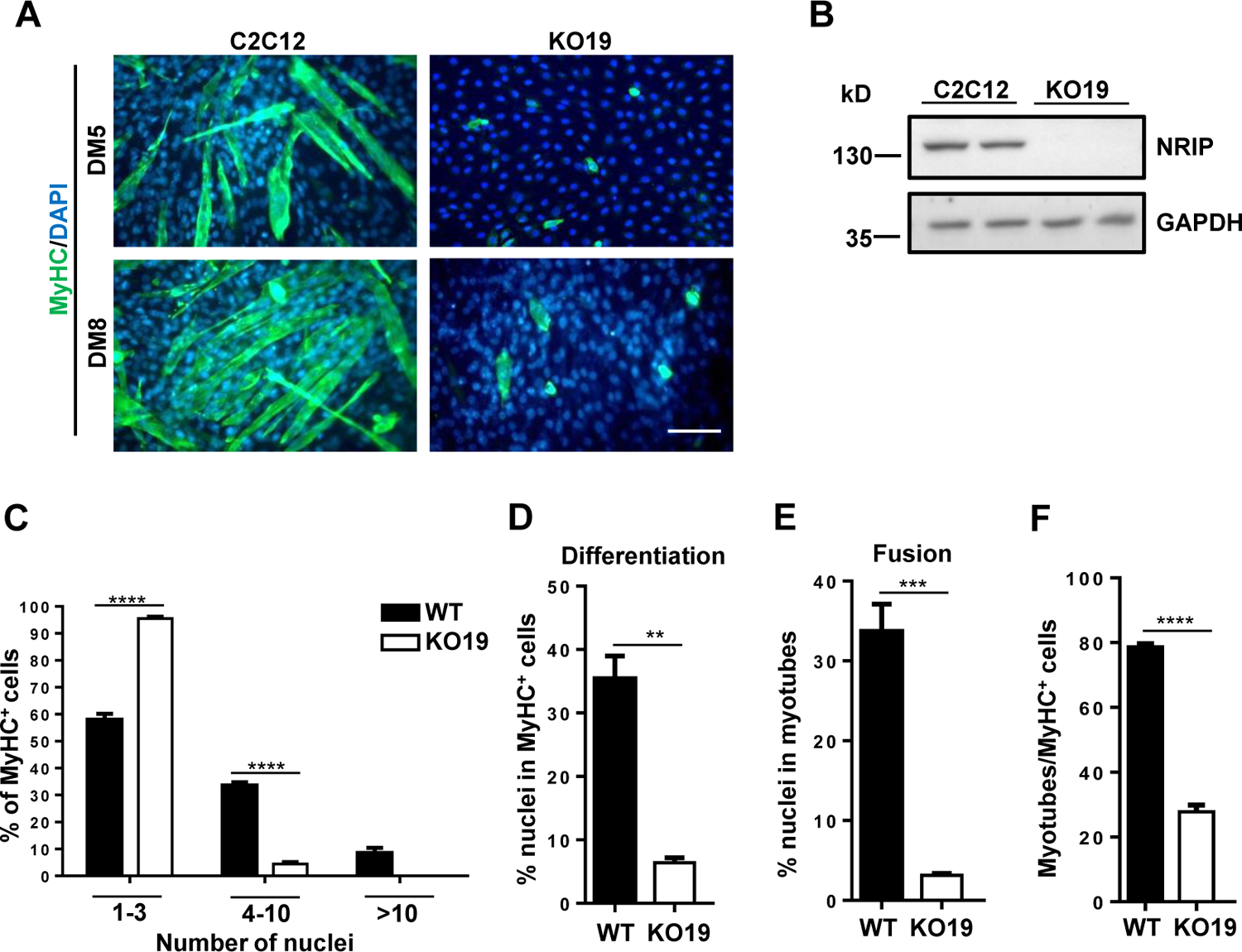
NRIP-null C2C12 cells (KO19 cells) reveal the decreased myotube formation. (A) Images of KO19 (NRIP-null C2C12 cells) for myotube formation. NRIP-null C2C12 cells were generated by CRISPR-Cas9 system. C2C12 and KO19 cells were differentiated for 5 (DM5) or 8 days (DM8) and stained with anti-MyHC antibody (green) and DAPI for nucleus (blue). Scale bar: 100 μm. (B) Western blot analysis of NRIP in KO19 cells. (C) The distribution of MyHC^+^ cells with 1-3 nuclei, 4-10 nuclei, and >10 nuclei at DM8 (N=3). (D) Differentiation index calculated as the percentage of nuclei in MyHC^+^ cells (including mononuclear and multinuclear cells) to total nuclei per field (N=3). (E) Fusion index measured by the percentage of nuclei contained (≥ 2 nuclei) of MyHC^+^ myotube to total nuclei per field (N=3). (F) The percentage of myotubes (≥ 2 nuclei in MyHC^+^ cells) to total MyHC^+^ cells (N=3). Data are mean ± SEM from 3 separate experiments. \*\**P* < 0.01, \*\*\**P* < 0.005, and \*\*\*\**P* < 0.001; student *t* test. See also Figure 3—figure supplement 1 and 2, Figure 3—source data 1 and 2. Source data 1: Numerical data for Figure 3C to 3F. Source data 2: Source WB blot for Figure 3B.

### Loss of NRIP reduces satellite cell activation and delayed regeneration in injured muscles

Most satellites are quiescent in mature skeletal muscles and will be activated in response to injury to initiate a regeneration program (De Bari et al., 2003) that includes satellite cell activation/proliferation (myoblast), myoblast differentiation into myocytes, and myocyte fusion for myotube formation. According to the Pax7 and MyoD expression patterns, differentiation during regeneration can be classified into three groups: Pax7^+^/MyoD^-^ cells (quiescent satellites), Pax7^+^/MyoD^+^ cells (myoblast activation/proliferation), and Pax7^-^/MyoD^+^ cells (myoblasts differentiating into myocytes) (Bentzinger et al., 2012; Zammit, Relaix, et al., 2006). In our previous study, the level of NRIP increased after muscle injury and was essential for skeletal muscle regeneration; however, its mechanisms were unclear (Chen et al., 2015). As shown in Figures 2 and 3, NRIP functions in myoblast differentiation and fusion. To identify the role of NRIP in satellite cell activation, we used Pax7 and MyoD expression markers. The tibialis anterior (TA) muscles of 8-week-old WT and cKO mice were injected with cardiotoxin (CTX) and harvested at 1, 3, and 6 days after treatment for immunofluorescence assay (Figure 3-figure supplement 2A and 2B). Numerical data for each time after CTX treatment are shown in Supplemental Figure 2C and 2D. On day 1 after injury, the proportion of Pax7^+^/MyoD^-^ cells was significantly lower in cKO than WT mice (29.38% vs. 37.88%) (Figure 3-figure supplement 2C and 2D), and that of Pax7^+^/MyoD^+^ cells was also lower (2.19% vs. 4.63%) (Figure 3-figure supplement 2C and 2D), which indicates that lack of NRIP reduces myoblast activation. On day 3 after injury, the proportion of Pax7^-^/MyoD^+^ cells was significantly lower in cKO mice than in WT mice (10.59% vs. 17.18%) (Figure 3-figure supplement 2C and 2D), which indicates that NRIP can promote myoblast differentiation into myocytes. However, on day 6, the proportion of Pax7^-^/MyoD^+^ cells was higher in cKO mice than in WT mice (17.75% vs. 11.35%) (Figure 3-figure supplement 2C and 2D), which indicates that NRIP deficiency delays the process of myogenesis differentiation. Thus, the myoblasts in WT mice began to activate/proliferate on day 1 and subsequently differentiated on day 3 after damage, but the cKO muscles showed delayed myoblast differentiation until day 6 after injury. This finding is consistent with our previous report, which found that NRIP deficiency delays muscle regeneration (Chen et al., 2015). Hence, NRIP affects satellite cell activation and differentiation during muscle regeneration in the skeletal muscles of mice.

### NRIP can rescue myotube formation in NRIP-null C2C12 cells

Several actin-binding proteins regulate cell fusion via actin polymerization/depolymerization during myotube formation (Berger et al., 2008; Nowak et al., 2009; Richardson et al., 2007). As shown in Figures 2 and 3, the loss of NRIP caused defective myoblast fusion. To examine whether NRIP could restore myotube formation in NRIP-null cells (KO19 cells), KO19 cells were overexpressed with NRIP. An immunofluorescence assay revealed that KO19/NRIP cells had larger, longer, and more myotubes than KO19/vector cells (Figure 4A), and NRIP was expressed in KO19/NRIP cells (Figure 4B). We calculated the proportion of myotubes in KO19/NRIP and KO19/vector cells containing 1–3 nuclei (93.71% vs. 98.75%, *P* < 0.01), 4-10 nuclei (5.72% vs. 1.25%, *P* < 0.01), and > 10 nuclei (0.57% vs. none) (Figure 4-figure supplement 1A). The differentiation was significantly higher in KO19/NRIP than in KO19/vector cells (5.72% vs. 3.06%, *P* < 0.05; Figure 4C, left panel), so re-expression of NRIP in NRIP-null KO19 cells could rescue differentiation. KO19/NRIP cells also showed increased ability of myoblast fusion as compared with KO19/vector cells (3.37% vs. 1.00%, *P* < 0.01, Figure 4C, right panel), which implies that re-expression of NRIP in KO19 cells can restore myotube formation. To demonstrate the independent NRIP role in myoblast fusion, the proportion of myotubes (MyHC^+^ cells ≥ 2 nuclei) to total MyHC^+^ cells was higher in KO19/NRIP than KO19/vector cells (32.13% vs. 16.28%, *P* < 0.001, Figure 4D). Collectively, NRIP can rescue myotube formation by promoting myoblast differentiation and fusion, and NRIP alone can also induce myoblast fusion.

**Figure 4.**
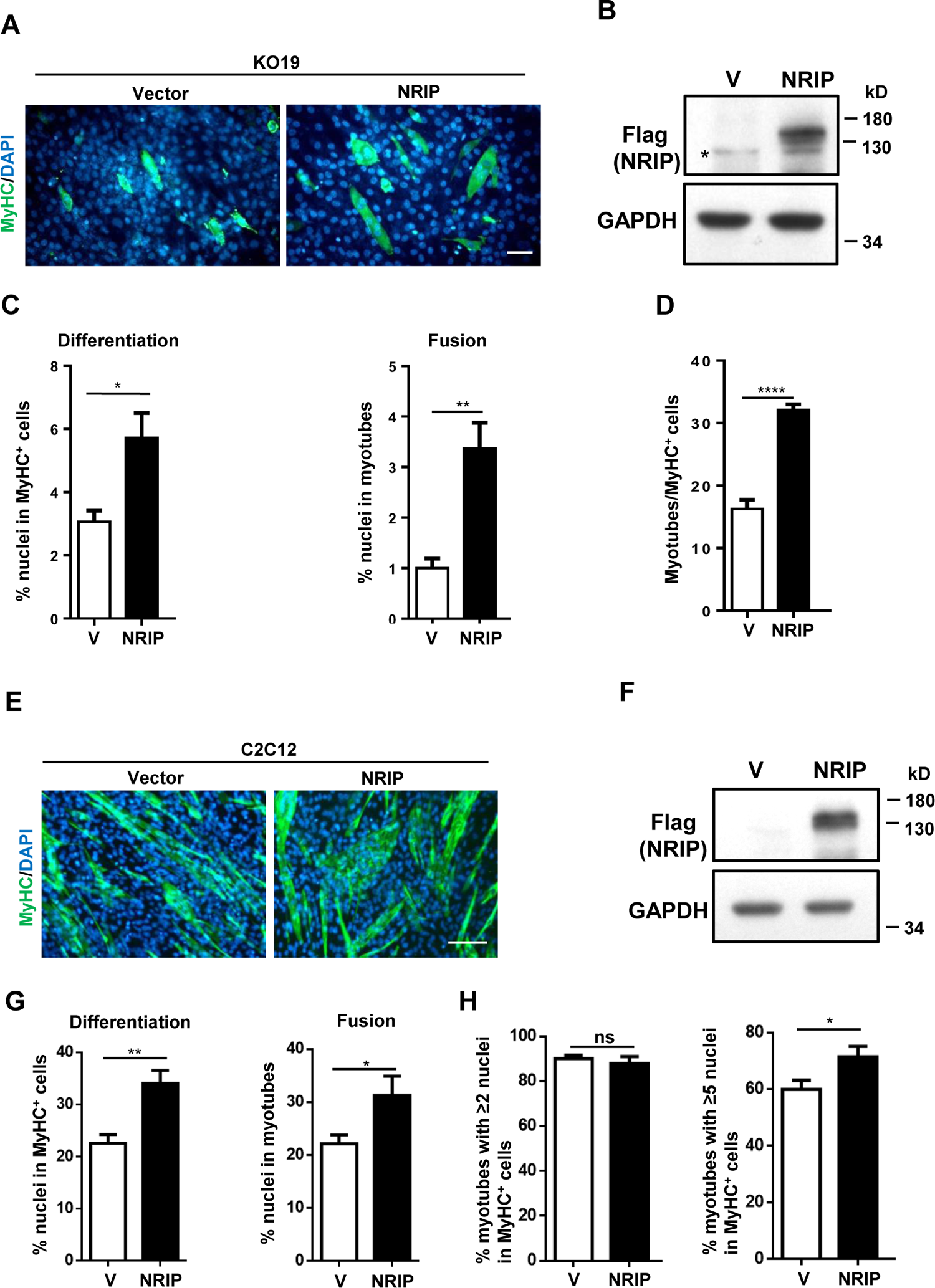
NRIP can rescue myotube formation in NRIP-null (KO19) and overexpression of NRIP in C2C12 increases the size of myotubes. (A) Images of overexpressing NRIP in NRIP-null cells can rescue myotube formation. KO19 cells were transfected with Flag vector or Flag-NRIP plasmid for 24 h and shifted to differentiation medium for 8 days, then stained with anti-MyHC antibody (green) and DAPI for nucleus (blue) to examine myotube formation. Scale bar: 100 μm. (B) Western blot analysis showed the protein expression of Flag-NRIP in KO19 cells. (C) Differentiation (left panel) and fusion (right panel) index (N=5). The method for counting differentiation and fusion was described in Figure 2 and 3. (D) The percentage of myotubes (≥ 2 nuclei in MyHC^+^ cell) to total MyHC^+^ cells (N=5). (E) Images of overexpressing NRIP in C2C12 can increase myotube size. C2C12 cells were transfected with Flag-NRIP plasmid for 24 h and shifted to differentiation medium for 5 days, then stained with anti-myosin heavy chain (MyHC) antibody (green) and DAPI for nucleus (blue). Scale bar: 100 μm. (F) Protein expression of NRIP in C2C12 cells. (G) Differentiation (left panel) and fusion (right panel) index (N=6). (H) Percentage of myotubes (≥ 2 nuclei in MyHC^+^ cells; left) or (≥ 5 nuclei; right) to total MyHC^+^ (N=6). Data are mean ± SEM from five (panels A-D) or six separate experiments (panels E-H). \**P* < 0.05, \*\**P* < 0.01, \*\*\*\**P* < 0.001 ns, no significance; student *t t*est. See also Figure 4—figure supplement 1, Figure 4—source data 1 and 2. Source data 1: Numerical data for Figure 4C, 4D, 4G and 4H. Source data 2: Source WB blot for Figure 4B and 4F.

### Overexpression of NRIP in C2C12 increases the multinucleated nuclei formation of myotubes

As shown in Figures 2 and 3, the deficiency of NRIP decreased multinucleated myotube formation. To further examine whether NRIP could contribute to large myotube formation, cells were transfected with Flag-NRIP plasmid and differentiated for 5 days (Figure 4E), and Flag-tagged NRIP could be detected in NRIP-containing C2C12 cells (Figure 4F). C2C12/NRIP incubation significantly contributed to multinucleated myotubes as compared with the control (C2C12/vector) (Figure 4-figure supplement 1B). Consistent with the effect of NRIP, the differentiation and fusion indexes in C2C12/NRIP were both increased; the differentiation index was higher for C2C12/NRIP than C2C12/vector (34.12% vs. 22.59%, *P* < 0.01, Figure 4G, left panel), as was the fusion index (31.24% vs. 22.16%, *P* < 0.05, Figure 4G, right panel). For the independent role of NRIP in myocyte fusion, the proportion of myotubes (≥ 2 nuclei per MyHC^+^ cell) to total MyHC^+^ cells was not significantly different with C2C12/NRIP and C2C12/vector cells (87.81% vs. 90.03%, *P* = 0.538, Figure 4H, left panel) but was different with ≥ 5 nuclei per MyHC^+^ cell (71.40% vs. 59.95%, *P* < 0.05, Figure 4H, right panel), which indicates that NRIP can enhance myoblast fusion to form large myotubes. Thus, NRIP can increase the formation of multinucleated myofibers.

### NRIP is a novel actin-binding protein

As shown in Figures 2 and 3, NRIP plays a role in myoblast fusion. Actin-binding proteins, such as Arp2/3 and Nap-1, are important for actin cytoskeleton remodeling during myoblast fusion (Berger et al., 2008; Dalkilic et al., 2006; Mattila & Lappalainen, 2008; Nowak et al., 2009; Richardson et al., 2007). NRIP directly interacts with α-actinin 2 (ACTN2), a well-known actin-binding protein, to facilitate F-actin bundling in cardiomyocytes (Yang et al., 2019). Hence, we hypothesized that NRIP might regulate myoblast fusion by interacting with actin. To identify whether NRIP directly interacted with actin *in vitro*, His-MBP and His-MBP-NRIP generated from *E. coli* [Figure 5A, (a)] were incubated with F-actin (from bovine cardiac muscle; AD99, cytoskeleton), respectively, and were subjected to immunoprecipitation and His-pull down assay. The results showed that NRIP interacted with F-actin by immunoprecipitation with anti-actin antibody [Figure 5A, (b)]. On the other hand, the F-actin was pull-downed with His-MBP-NRIP using Ni-NTA beads [Figure 5A, (c)]. In sum, NRIP can reciprocally bind to F-actin *in vitro.* To further confirm *in vitro* binding, His-MBP-NRIP was incubated with F-actin, and a low-speed sedimentation assay was performed (Yang et al., 2019). The result revealed that the F-actin pellets increased when His-MBP-NRIP was incubated with F-actin (Figure 5B, lane 8) as compared to when His-MBP (control) was incubated with F-actin (Figure 5B, lane 6). Quantitation analysis revealed that NRIP significantly facilitated F-actin bundling (Figure 5B, right). To clarify the interaction of NRIP and actin in mammalian cells, we used an immunoprecipitation assay with Flag-NRIP and mCherry-actin plasmids co-transfected into 293T cells. NRIP could be precipitated by mCherry-actin (Figure 5C, left panel lane 4), and actin could be detected by the immunoprecipitation of Flag-tagged NRIP (Figure 5C, right panel lane 4). Thus, NRIP is a novel actin-binding protein that can interact with actin both *in vitro* and *in vivo*.

**Figure 5.**
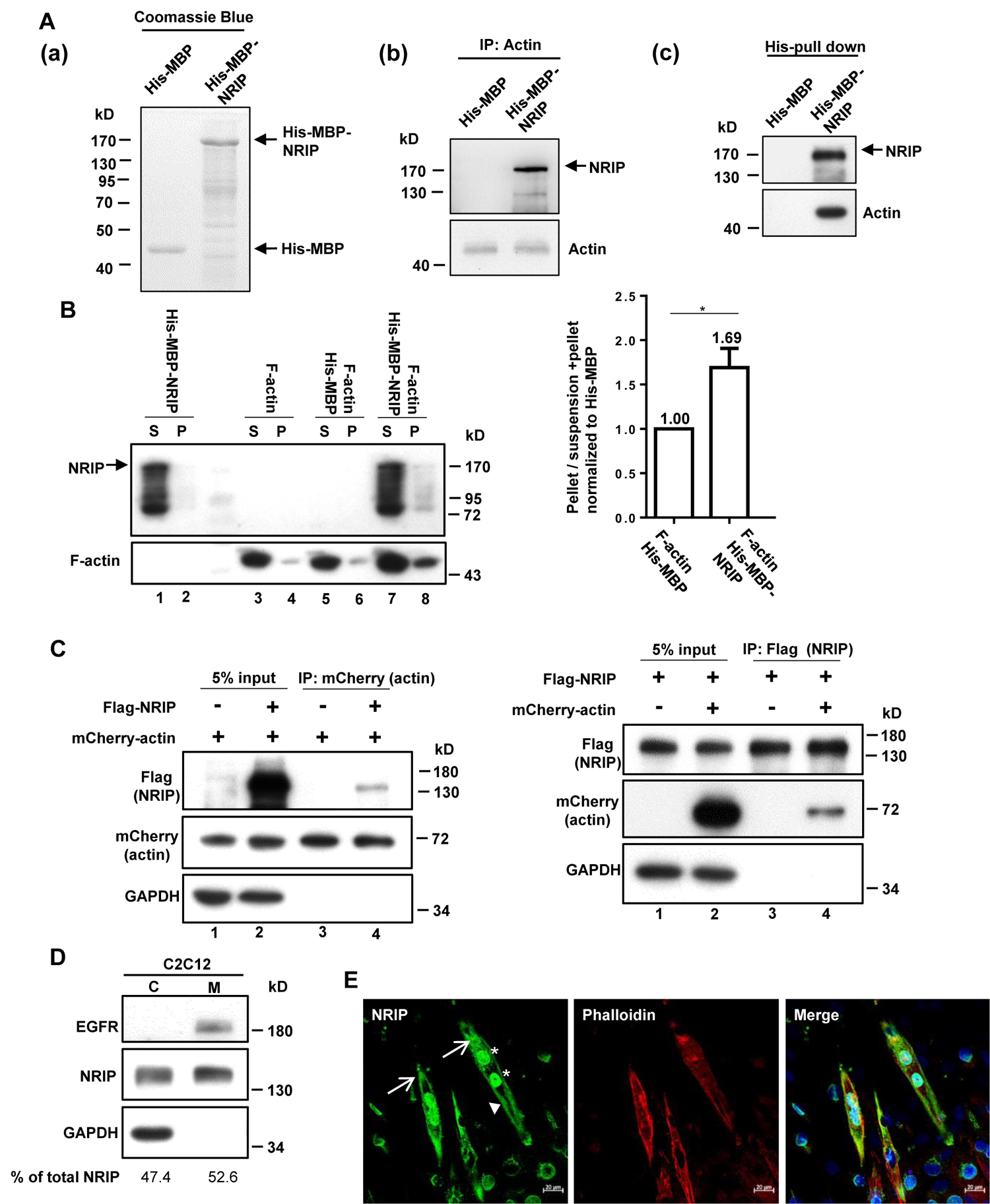
NRIP can directly and reciprocally interact with actin as a novel actin-binding protein. (A) NRIP directly interacts with F-actin *in vitro*. (A (a)) The expression proteins of His-MBP and His-MBP-NRIP from bacteria were purified separately; and examined by Coomassie blue staining. The arrows indicate His-MBP and His-MBP-NRIP. (A (b)) The His-MBP-NRIP from bacteria were incubated with F-actin from bovine cardiac muscle (AD99, Cytoskeleton). The protein mixtures were immunoprecipitated (IP) with anti-actin antibody and analyzed by western blot with anti-NRIP. (A (c)) The His pull-down was performed by Ni-NTA agarose and immunoblotted by anti-actin. (B) NRIP interacts with F-actin *in vitro* using low-speed sedimentation. Purified His-MBP and His-MBP-NRIP from bacteria were incubated with bovine cardiac muscle actin respectively and centrifuged by low-speed sedimentation to separate supernatant (S) and pellet fractions (P). Left panel: lanes 1 and 2 for His-MBP-NRIP only; lanes 3 and 4 for F-actin only; lanes 5 and 6 for His-MBP with F-actin; lane 7 and 8 for His-MBP-NRIP with F-actin. Right: quantitation of the ratio of sedimented actin (pellet/total) from three independent experiments. Data are mean ± SEM from 3 separate experiments. \**P* < 0.05; student *t t*est. (C) NRIP interacts with F-actin *in vivo*. 293T cells were transfected with expression vectors for Flag-tagged NRIP or control vector, along with either mCherry-tagged F-actin expression plasmid or mCherry vector. Cell extracts (1mg for each group) were immunoprecipitated (IP) with anti-mCherry (actin) (left panel) or with anti-Flag (NRIP) (right panel) antibody and immunoblotted with indicated antibodies. GAPDH as a loading control. (D) Subcellular localization of NRIP in C2C12 cells. C2C12 myoblasts were harvested; cytosolic (C) and membrane (M) fractions were extracted as described in Materials and methods. EGFR: epidermal growth factor receptor as a positive control of membrane protein, GAPDH: as a positive control of cytosolic proteins. % of total NRIP was counted by the intensity ratio of cytosolic or membrane NRIP to total NRIP those were measured by ImageJ software. Data are representative of three independent experiments. (E) Endogenous NRIP locates at plasma membrane, nucleus and cytoplasm. C2C12 myoblasts were differentiated for 4 days for subcellular location analysis using immunofluorescence staining of NRIP (green), phalloidin for cell membrane (red, F-actin staining, arrowhead), and DAPI for nucleus (blue, asterisk). The cytoplasm location shown by arrow. Scale bar: 20 μm. See also Figure 5—source data 1 to 4. Source data 1: Numerical data for Figure 5B. Source data 2: Source WB blot for Figure 5A and 5B. Source data 3: Source WB blot for Figure 5C-left panel. Source data 4: Source WB blot for Figure 5C-right panel and 5D.

### NRIP is located at the plasma membrane, nucleus, and cytoplasm in C2C12 cells

Apart from actin-binding proteins in regulating myoblast fusion, several of the membrane-bound proteins that participate in cell fusion include myomaker, myomixer (also named myomerger and minion), and stabilin-2 (Bi et al., 2017; Millay et al., 2013; Park et al., 2016; Quinn et al., 2017; Zhang et al., 2017). Thus, we examined the subcellular localization of NRIP in C2C12 cells by fractionating C2C12 myoblasts into cytosolic and membrane fractions. NRIP was localized in both the cytoplasm and cell membrane at nearly equal proportions (47.4% and 52.6%, respectively; Figure 5D). Additionally, an immunofluorescence staining of C2C12 cells showed that NRIP colocalized with actin and was located at the membrane (co-stained with phalloidin, an F-actin binding protein); NRIP was also located at the nucleus (colocalized with DAPI) or cytoplasm (Figure 5E). This finding is consistent with our previous studies showing that NRIP is a nucleus protein and functions as a transcription cofactor to mediate androgen receptors to activate gene expression (Chen et al., 2008; Tsai et al., 2005). In addition, NRIP is located in cytoplasm to bind with CaM or ACTN2 of sarcomere (Chen et al., 2015; Yang et al., 2019). Here, we found NRIP co-localizing with actin at the cell membrane. Thus, NRIP is an actin-binding protein located at the cell membrane, cytosol, and nucleus.

### NRIP is a novel invadosome protein for myoblast fusion

As shown in Figures 2 to 4, NRIP was required for myoblast fusion and directly interacted with F-actin (Figure 5). Invadosomes are F-actin–enriched protrusions that are important for invasion and fusion pore formation (Chen, 2011; Chuang et al., 2019). The enriched F-actin with Tks5 (invadosome scaffold protein) and cortactin (actin binding protein as an invadosome marker) in C2C12 myotubes can form an invadosome that is required for myoblast fusion (Chuang et al., 2019; Deng et al., 2015; O’Connell et al., 2019). Hence, we hypothesized that NRIP might be involved in invadosome formation through actin binding to regulate myoblast fusion. We first examined the expression profiles of NRIP, Tks5, and cortactin during C2C12 myoblast differentiation. The NRIP levels were in compliance with expression patterns of Tks5 and Cortactin protein (Figure 6A), which are the gradually increasing expression upon myoblast differentiation (Chuang et al., 2019) and the increasing myosin heavy chain (MyHC) expression (Figure 6A). The immunofluorescence results also showed that NRIP was colocalized with cortactin (Figure 6B, upper), Tks5 (Figure 6B, middle), and F-actin (Figure 6B, lower), respectively, at the tips of C2C12 (Figure 6B; arrows). This indicates that NRIP expression is enriched in invadosomes during C2C12 myoblast differentiation. To further confirm whether NRIP is required for invadosome formation, we transfected pSuper-shNRIP into C2C12 cells to knockdown the expression of endogenous NRIP. The expression of NRIP was significantly decreased in NRIP-knockdowned C2C12 cells compared to control cells by the immunofluorescence stain (Figure 6C) and western blot analysis (Figure 6D), respectively. Moreover, the knockdown of NRIP resulted in the dispersed expression of F-actin in C2C12 myoblasts (Figure 6C) without affecting the protein expression of F-actin (Figure 6D). This implies that NRIP is a novel component of invadosomes. Furthermore, we used a Lifeact-RFP labeling assay to distinguish the expression pattern of NRIP in attacking cells and receiving cells during myoblast fusion. The Lifeact-RFP is a 17-amino-acid peptide derived from the actin-binding protein 140 (Abp140) of *S. cerevisiae* (Riedl et al., 2008) and specifically binds with F-actin, which is strongly expressed in attacking cells during myoblast fusion (Chuang et al., 2019). Hence, the Lifeact-RFP was transfected into C2C12 cells to label F-actin; the result showed NRIP expression was enriched in Lifeact-RFP expressed attacking cells and was dispersed in receiving cells between the two close-positioned, 3-day differentiated myoblasts (Figure 6E). This indicates that NRIP is required for invadosome formation through actin binding.

**Figure 6.**
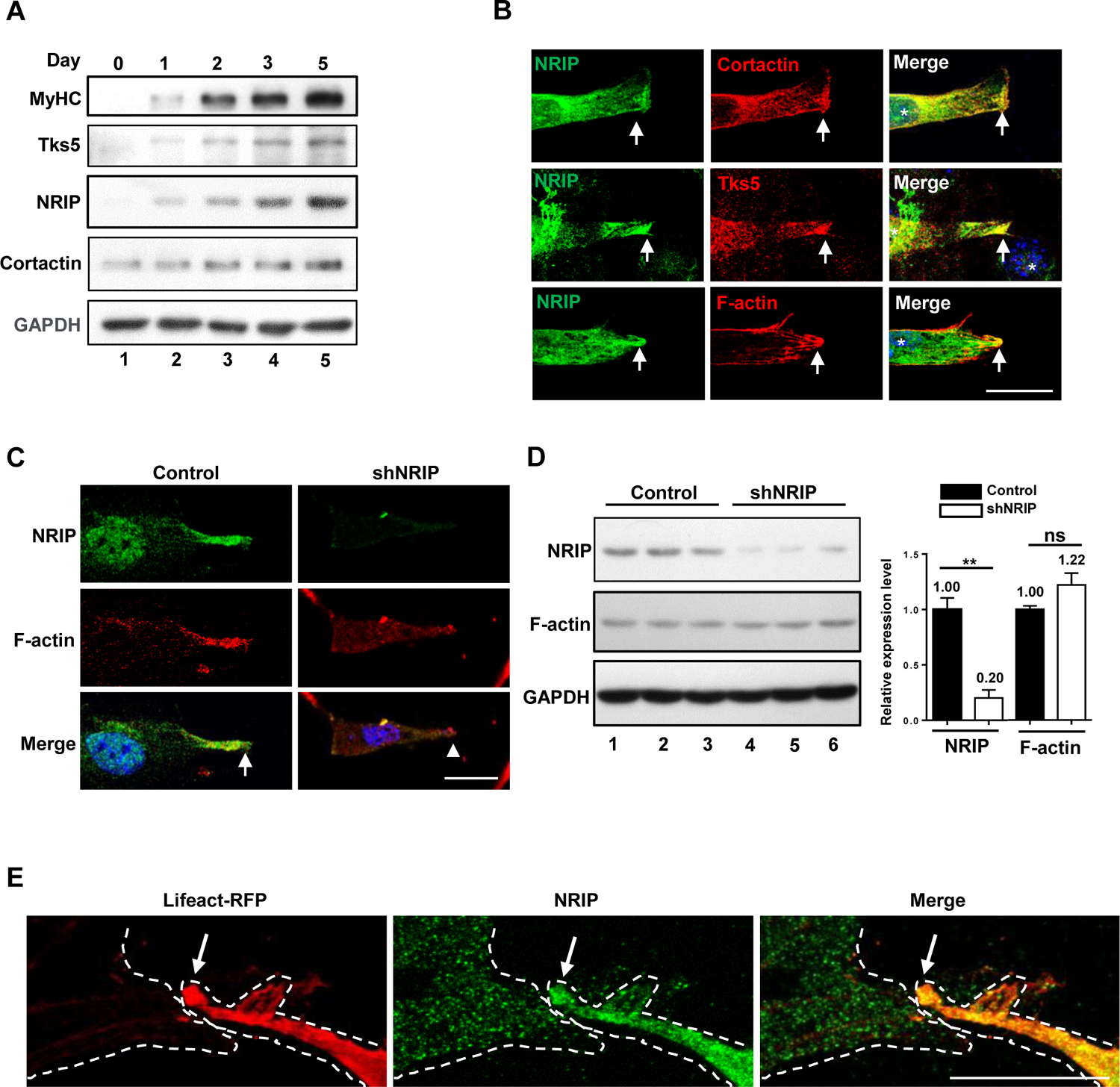
NRIP is a novel invadosome protein for myoblasts fusion. (A) The expression profiles of NRIP and invadosome proteins (Tks5, invadosome scaffold protein), cortactin (enriched in invadosome and bound with actin) during myoblast (C2C12) differentiation. MHC (myosin heavy chain) as a differentiation marker. The GAPDH as a loading control. (B) Endogenous NRIP locates at tips of invadosome sites during cell-cell fusion. The enriched NRIP colocalized with cortactin (upper), Tks5 (middle), and F-actin (lower) at 3 days of C2C12 differentiation. The arrows indicate the tip of cells. Green: NRIP; red: each component of invadosome; blue: DAPI for nucleus counter stain. The arrows indicate the tip of cells. Scale bar: 100 μm. (C) The knockdown NRIP expression reduces the tip distribution of F-actin. The cells were transfected with pSuper-shNRIP plasmid for 24 h and shifted to differentiation medium for 5 days, then stained with anti-F-actin (red) and anti-NRIP antibodies (green) and DAPI for nucleus (blue). Scale bar: 100 μm. Arrow: the enriched NRIP and F-actin expression at tips of C2C12 cells. Arrowhead: the dispersed F-actin at tip of NRIP knockdown C2C12 cells. Scale bar: 100 μm. (D) The F-actin protein expression is comparable between NRIP knockdown cells and wild type. The left panel showed expression of NRIP was decreased in NRIP-knockdown C2C12 cells compared to C2C12 cells at differentiated day 5; but the expression of F-actin was not change. Right: quantification of NRIP and F-actin protein expression from left panel (N=3). Data are mean ± SEM from 3 separate experiments. \*\**P* < 0.01; ns, no significance; student *t t*est. The GAPDH as a loading control. (E) The invadosome containing NRIP and F-actin distributes asymmetry during myoblast fusion. The cells transfected with Lifeact-RFP (stained for F-actin located at cell membrane) at 3 days of C2C12 cell differentiation. The dashed lines represent the cell periphery of two cells (one represents receiving cell, the other is attacking cell containing invadosome). In attacking cells, the endogenous NRIP (green, middle) was strongly expressed with F-actin located at cell membrane (left); but in receiving cells, endogenous NRIP was evenly distributed in cells (middle). Arrows (attacking cell) indicate enriched F-actin at cell membrane (red, left); and NRIP in middle (green). Scale bar: 100 μm. See also Figure 6—source data 1 and 2. Source data 1: Numerical data for Figure 6D. Source data 2: Source WB blot for Figure 6A and 6D.

### NRIP participates in invadosome protrusion during myoblast fusion

Myoblast fusion can be divided into several steps, including migration, recognition, adhesion, invasion, and fusion (Abmayr & Pavlath, 2012). For invasion, the asymmetrical distribution of the invadosome is constructed between two fusion partners in *Drosophila* embryo and murine myoblast C2C12 cells (Abmayr & Pavlath, 2012; Chuang et al., 2019). We found that NRIP was a novel actin-binding protein (Figure 5) that could also be an invadosome protein (Figure 6). To further demonstrate whether NRIP participated in invadosome protrusion, we used time-lapse microscopy to analyze the dynamics of NRIP expression in C2C12 myoblasts during myoblast fusion. C2C12 cells were co-transfected with EGFP-NRIP and mCherry-actin for 24 h and shifted to the differentiation medium for another 24 h. The immunofluorescence stain showed that EGFP-NRIP was colocalized with mCherry-actin to form the invadosome of C2C12 (Figure 7A, arrows). To observe live cell imaging, cells were replated on matrigel-coated cover glass and subjected to time-lapse microscopy. Live cell imaging showed EGFP-NRIP and mCherry-actin were concentrated to form foci in the cytoplasm of C2C12 cells (0 min); then, the foci protruded toward the cell membrane to form a protrusive invadosome (30 min) (Figure 7B and Figure 7-figure supplement video 1). The protrusive invadosome elongated and moved closer to the receiving cell (60 min) (Figure 7B and Figure 7-figure supplement video 1). Finally, the protrusive invadosome made contact the membrane of the receiving cell, and the EGFP-NRIP and mCherry-actin were diffused to the cytoplasm of the receiving cell (90 min) (Figure 7B and Figure 7-figure supplement video 1). In summary, NRIP formed a focus and colocalized with F-actin to produce an invadosome that elongated over the time of fusion, which further supported the idea that NRIP interacts with F-actin to form an invadosome for myoblast fusion.

**Figure 7.**
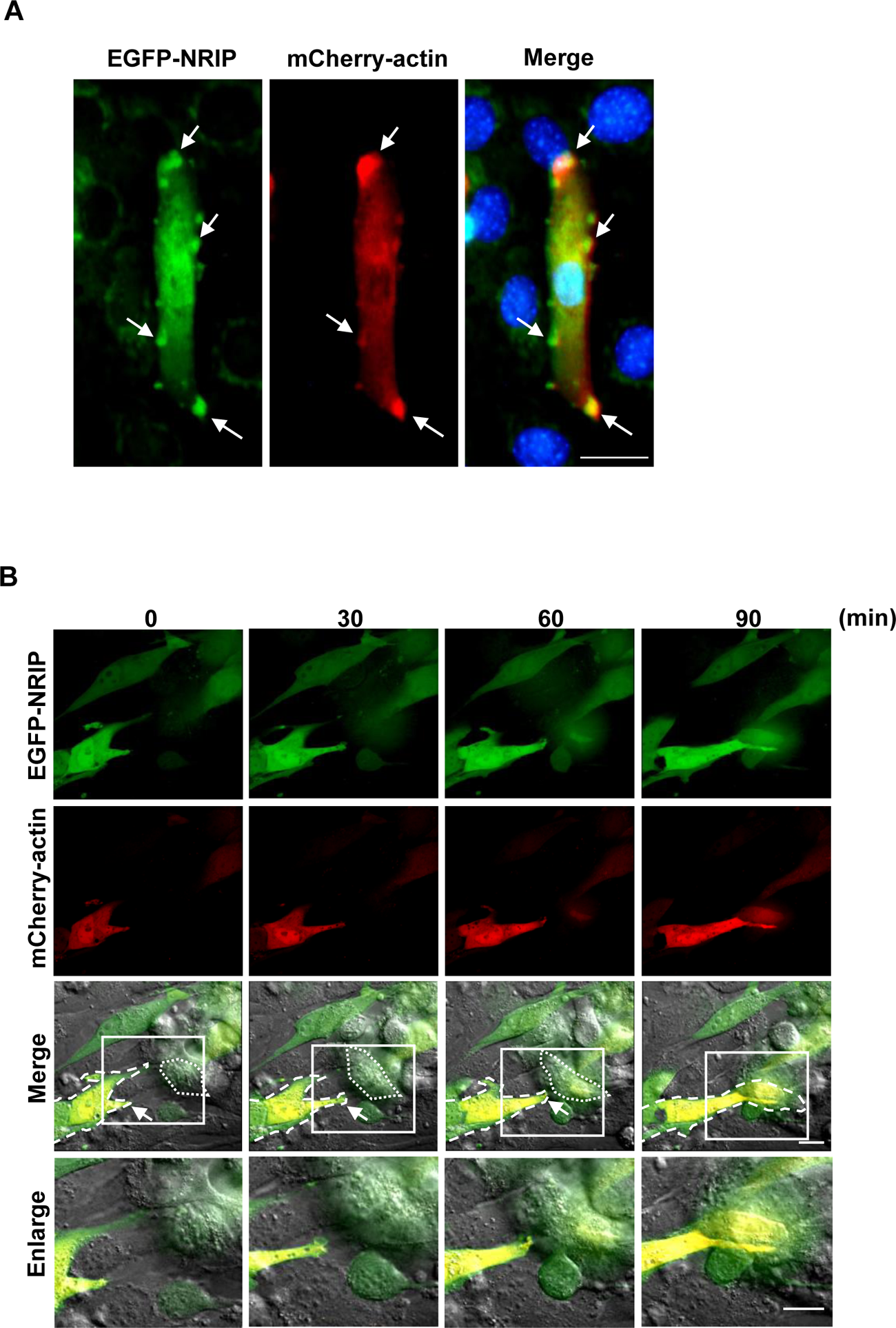
The dynamics of NRIP expression in actin-enriched invadosome during myoblasts fusion. (A) EGFP-NRIP and mCherry-actin-based finger-like structures in differentiated C2C12 cells. The Z-stack of merge image showed EGFP-NRIP and mCherry-actin were colocalized at several invadosomes of C2C12 myotubes indicated by the arrows. DAPI (blue) for nuclear counter stain. Scale bar: 100 μm. (B) The dynamics of NRIP expression along with the invadosome protrusion during myoblast fusion. C2C12 cells were co-transfected with mCherry-actin (red) and EGFP-NRIP (green). After differentiation for 3 days, cells were then transferred into matrigel-coated plates for monitoring the process of myoblast fusion using time-lapse microscopy. Initially, the invadosome containing mCherry-actin (red) and EGFP-NRIP (green) located at the cell tip of attacking cells (0 min). White arrows of merged images indicated the expression of EGFP-NRIP along with mCherry-actin in invadosome structure of attacking cells. Boxed regions were enlarged and shown in Enlarge panel. At 30 min, the invadosome elongated from attacking cells (dash line) toward the receiving cells (dot line). At 60 min, the invadosome stretched; and at 90 min, the invadosome contacted to the receiving cells and fusion occurred between attacking cells and receiving cell. Scale bar: 20 μm. See also Figure 7—figure supplement video 1.

### NRIP participates in cell–cell fusion

NRIP-deficient mice show a high proportion of small myofibers during muscle regeneration (Chen et al., 2015). As shown in Figures 5 to 7, NRIP is an actin-binding protein that promotes invadosome protrusion during myoblast fusion. To further demonstrate NRIP’s participation in cell–cell fusion, we transfected C2C12 cells with an mCherry vector, which revealed a red color. KO19 cells (NRIP-null C2C12) were then transfected with an EGFP vector as a control or EGFP-NRIP as an experimental group, which yielded a green color. Then, C2C12 and KO19 cells were mixed for differentiation for myotube formation (Figure 8A). Only EGFP^+^ (green) or mCherry^+^ (red) cells indicated non-fusion with the other kinds of cells, and co-positive cells represented the fusion that occurred between the two kinds of cells. The KO19/EGFP-NRIP+C2C12/mCherry cells showed larger fused myotubes with more myonuclei (co-positive for green and red, marked by arrows; Figure 8B) as compared with the control (KO19/EGFP+C2C12/mCherry). We calculated the number of co-positive myotubes with ≥ 3 nuclei to the total number of co-positive myotubes; KO19/EGFP-NRIP+C2C12/mCherry cells showed larger co-positive myotubes than the controls (control KO19/EGFP+C2C12/mCherry set as 1, 1.23 for KO19/EGFP-NRIP+C2C12/mCherry, *P* < 0.05; Figure 8C). Thus, NRIP participates in cell–cell fusion to increase the size of myotubes.

**Figure 8.**
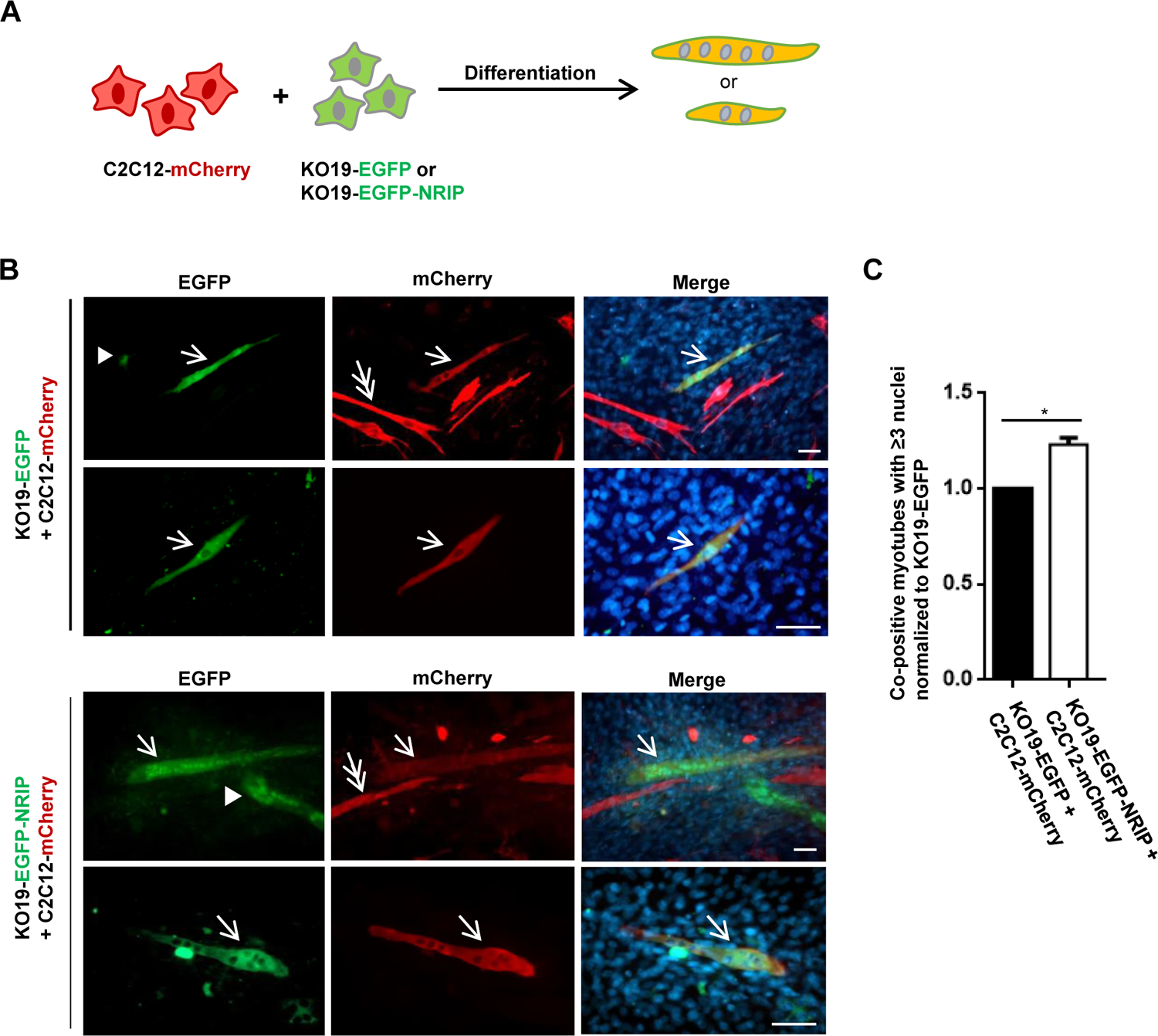
NRIP directly participates in myoblast fusion. (A) The schematic illustration of cell fusion assay. C2C12 cells were transfected with mCherry vector to show red color; and KO19 cells (NRIP-null) were transfected with EGFP vector as a control group or EGFP-NRIP as an experimental group, and displayed as a green color, respectively. Then, C2C12 and KO19 were mixed together in a ratio of 1:1 (cell density 7×10^4^ cells/cm^2^ for each cell line) and shifted to differentiation medium for 12 days to form myotubes. (B) The images of cell–cell fusion. The mixed cells were stained with anti-EGFP (green) and anti-DsRed for mCherry (red) to enhance the fluorescence intensity. Single green (arrowhead) or single red (double arrow) means no fusion. Co-positive cells, green and red (arrow), indicate the fused myotubes that express both EGFP and mCherry at the same time. Scale bar: 100 μm. (C) Quantitative analysis. The number of co-positive myotubes (≥ 3 nuclei) to total number of co-positive myotubes (N=3). Data are mean ± SEM from 3 separate experiments. \**P* < 0.05; student *t* test. See also Figure 8—source data 1. Source data 1: Numerical data for Figure 8C.

### Either NRIP WD6/7 or the IQ domain is responsible for actin binding, resulting in invadosome formation

NRIP could be involved in invadosome formation through actin binding for myoblast fusion (Figure 5 to 8). We further investigated which domain of NRIP was responsible for actin binding and determined the relationship between actin binding and invadosome formation. We used NRIP-FL (full length), NRIP-C (C fragment containing two WD40 domains and one IQ motif), NRIPΔIQ (full length without IQ motif), C-ΔWD6/7 (C fragment without WD40 domains but containing one IQ motif), C-ΔWD6/7ΔIQ (C fragment without WD40 domains and IQ motifs), and NRIP-WD6/7 (the sixth and seventh WD40 domains in NRIP-C fragment) to map the domain of NRIP for actin binding in 293T cells (Figure 9A). Immunoprecipitation assay revealed that NRIP-FL, NRIP-C, NRIPΔIQ, and C-ΔWD6/7 could bind to actin (Figure 9B, lanes 2-5). NRIPΔIQ, containing seven WD40 domains, could interact with actin (Figure 9B, lane 4), which suggests that the WD40 domain may be responsible for actin binding. Also, C-ΔWD6/7, with only one IQ motif, could bind with actin (Figure 9B, lane 5), so the binding might be through the IQ motif to indirectly bind with ACTN2 because NRIP can interact with the EF-hand of ACTN2 through its IQ motif (Yang et al., 2019) and ACTN2 is known as an actin-binding protein (Mimura & Asano, 1986). To further clarify the WD40 domain and IQ motif for actin binding, we constructed the C-ΔWD6/7ΔIQ plasmid (lack of WD40 domains and IQ domain), with lost actin binding (Figure 9C, lane 2) and NRIP-WD6/7 that could interact with actin (Figure 9C, lane 4); this indicates that the NRIP WD6/7 domain is responsible for actin binding. To further confirm the direct binding between the NRIP WD40 domain and actin, an *in vitro* His-pull down was performed. NRIP mutant proteins were generated from *E. coli* and represented by the Coomassie blue stain (Figure 9D, lower panel). NRIP mutant proteins were incubated with F-actin from bovine cardiac muscles (AD99, cytoskeleton) and pull-downed by nickel-charged affinity resin for subsequent WB analysis. The results of western blot showed that actin could be pull-downed with NRIP containing WD40 (NRIP-FL, NRIP-C, NRIPΔIQ, NRIP-WD6/7) (Figure 9D, lanes 2, 3, 4, and 7). In contrast, NRIP without WD6/7 domain (C-ΔWD6/7 and C-ΔWD6/7ΔIQ) lost the binding of actin (Figure 9D, lanes 5 and 6). This further supported the idea that NRIP with the IQ domain but no WD40 domain (C-ΔWD6/7) indirectly interacts with actin (shown in Figure 9B) was due to the interaction of the IQ domain with ACTN2 in C2C12 cells. Collectively, NRIP directly interacts with actin through its WD40 at the sixth and seventh domains (Figure 9D). To further investigate the correlation of the NRIP-actin interaction with invadosome formation, the expression and distribution of NRIP mutants and F-actin in differentiated C2C12 were examined by immunofluorescence staining (Figure 9E). The results showed that the immunofluorescence intensity of C-ΔWD6/7ΔIQ (i.e., the loss of actin binding) in actin-enriched invadosome was significantly decreased in C2C12 myotubes compared to NRIP-FL (0.91 vs. 2.59, *P* < 0.05, Figure 9E and 9F). The other mutants (such as NRIP-C, NRIPΔIQ, C-ΔWD6/7 and NRIP-WD6/7) having actin binding ability showed similar invadosome distribution as NRIP-FL (Figure 9E and 9F). Collectively, NRIP can be highly expressed at invadosome locations by interacting with actin either through WD40 domains for direct binding or indirectly through the IQ domain for ACTN2 to bind with actin.

**Figure 9.**
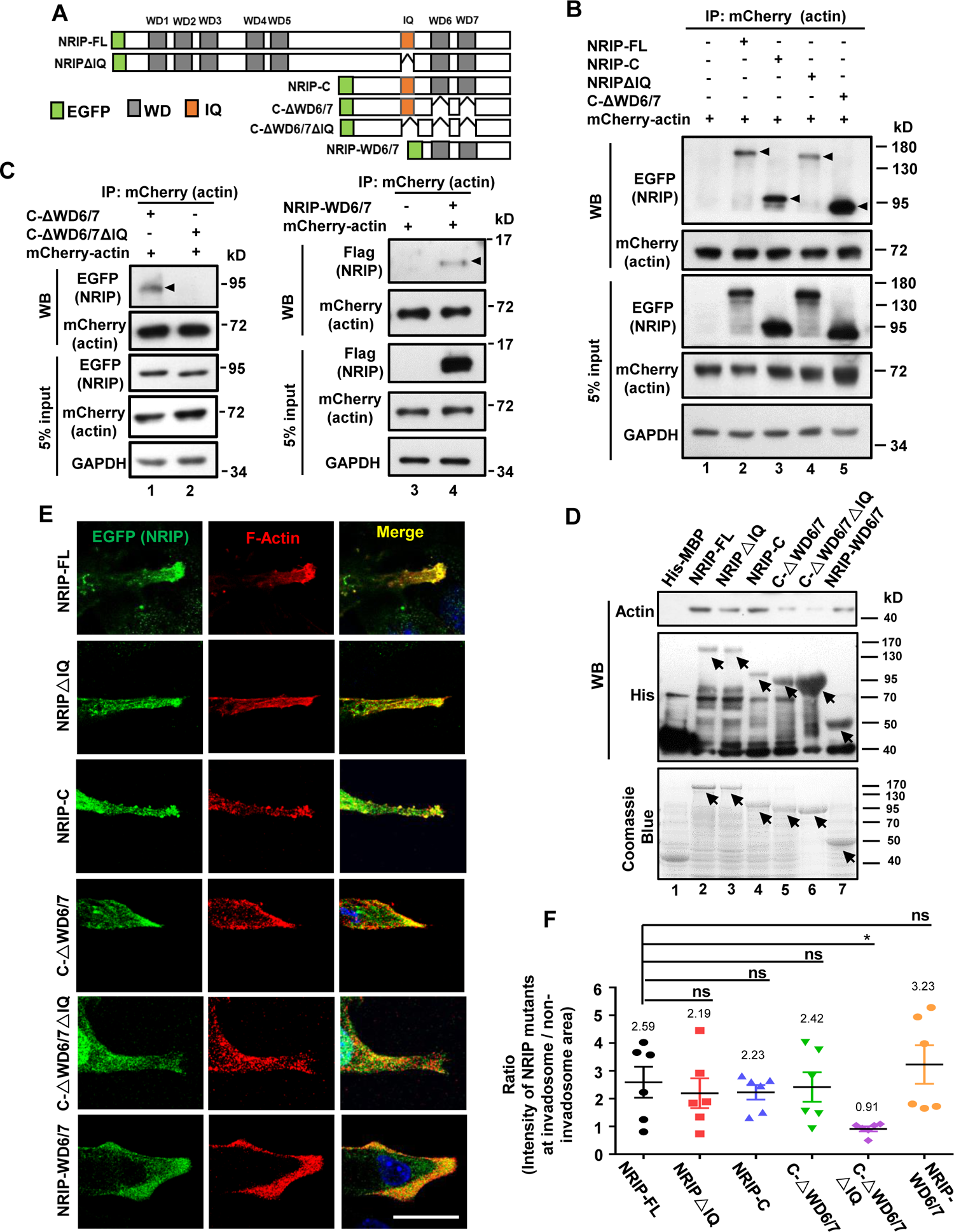
The correlation of NRIP-actin binding with invadosome formation using NRIP mutant analysis. (A) Schematic illustration of EGFP-tagged NRIP and NRIP deletion mutants. NRIP-FL (Full length), NRIPΔIQ (NRIP full length without IQ motif), NRIP-C (C fragment containing WD6/7 domains and one IQ motif), C-ΔWD6/7 (C-fragment containing one IQ motif and lack WD6/7), C-ΔWD6/7ΔIQ (C-fragment without WD6/7 domains and IQ domain) and NRIP-WD6/7 (only WD6/7 fragment). (B) Domain mapping of the NRIP-actin interaction in cells. 293T cells were transiently co-transfected with each NRIP truncated mutant and mCherry-actin plasmid for immunoprecipitation assay with anti-mCherry antibody for detecting mCherry-actin. Black arrowheads indicate the precipitated NRIP by actin (lane 2 to lane 5). GAPDH as a loading control. It indicates IQ domain and WD6/7 of C fragment in cells responsible for actin binding. (C) Either WD6/7 domains or IQ domain of NRIP can interact with actin in cells. The immunoprecipitated with actin showed on C-ΔWD6/7 (containing IQ domain, lack WD6/7 domains; lane 1) and NRIP-WD6/7 (containing WD6/7 domains; lane 4); but not C-ΔWD6/7ΔIQ (without IQ and WD6/7 domains). (D) NRIP-WD6/7 domains responsible for actin direct binding using *in vitro* His-pull down assay. The protein expression of His-MBP and His-MBP-NRIP mutants (NRIP-FL, NRIPΔIQ, NRIP-C, C-ΔWD6/7, C-ΔWD6/7ΔIQ and NRIP-WD6/7) from *E. coli* were examined by Coomassie blue staining (lower panel). The arrows indicate His-MBP and His-MBP-NRIP mutants’ proteins. His-MBP-NRIP mutant proteins from bacteria incubated with F-actin from bovine cardiac muscle (AD99, Cytoskeleton). The protein mixtures were pull-downed by Ni-NTA and immunoblotted with anti-actin antibody (WB: upper panel) and anti-His antibody (WB: lower panel). The NRIP-actin binding was absent in NRIP without WD6/7domain (C-ΔWD6/7, lane 5; C-ΔWD6/7ΔIQ; lane 6). In sum, IQ domain only could not bind with actin *in vitro* (C-ΔWD6/7; lane, 5); but WD6/7 domain could (WD6/7; lane 7). (E) NRIP localized at invadosome through actin interaction. The images showed expression of NRIP mutants (green) with endogenous F-actin (red) at invadosome. C2C12 myoblasts were transfected with EFGP-NRIP mutants and differentiated for 3 days. The C2C12 myotubes were stained with anti-EGFP (NRIP) and anti-F-actin antibodies; and imaged with confocal microscopy. The deficient actin binding of NRIP mutant (C-ΔWD6/7ΔIQ) reduced the enriched localization at invadosome. All actin binding NRIP mutants were significantly expressed at F-actin-enriched invadosome. DPAI (blue) for nuclear stain. (F) Quantification of NRIP mutants’ expression at invadosome from panel E (N=6). The invadosome was measured the enrichment of NRIP with F-actin; hence the ratio was quantified by the intensity of NRIP in F-actin focus divided by the intensity outside the invadosome. The six cells of each mutant were analyzed. Scale bars, 20 μm. Data are mean ± SEM from 6 separate experiments. \**P* < 0.05; student *t* test. See also Figure 9—source data 1 to 4. Source data 1: Numerical data for Figure 9F. Source data 2: Source WB blot for Figure 9B. Source data 3: Source WB blot for Figure 9C. Source data 4: Source WB blot for Figure 9D.

### Correlation between NRIP–actin binding and myotube formation

To explore the contribution of NRIP domains in myoblast fusion, we first examined whether NRIP affected myosin heavy chain (MyHC) expression. Western blot analysis revealed that NRIP and its mutants did not perturb MyHC expression of C2C12 myoblasts, as comparable amounts of MyHC protein were detected (Figure 10-figure supplement 1). Furthermore, to correlate NRIP-actin binding and myotube formation to the proportion of myotubes (≥ 5 nuclei per MyHC^+^ cell; shown in Figure 4H, right panel) for each mutant in C2C12 cells, the immunofluorescence staining of MyHC (Figure 10A) was investigated. The quantified data showed that loss of actin binding (C-ΔWD6/7ΔIQ mutant cells) significantly decreased for multinucleated myotube formation in comparison to NRIP-FL (46.77% vs. 65.18%, *P* < 0.01, Figure 10B); the other NRIP mutants (NRIP-C, NRIPΔIQ, C-ΔWD6/7 and NRIP-WD6/7) that had actin-binding abilities exhibited similar myotube formation as NRIP-FL (Figsure 10A and 10B). Collectively, NRIP is a novel actin-binding protein via the WD40 domain or the IQ motif for interaction, and actin binding is correlated with invadosome formation and myoblast fusion (Figure 10C).

**Figure 10.**
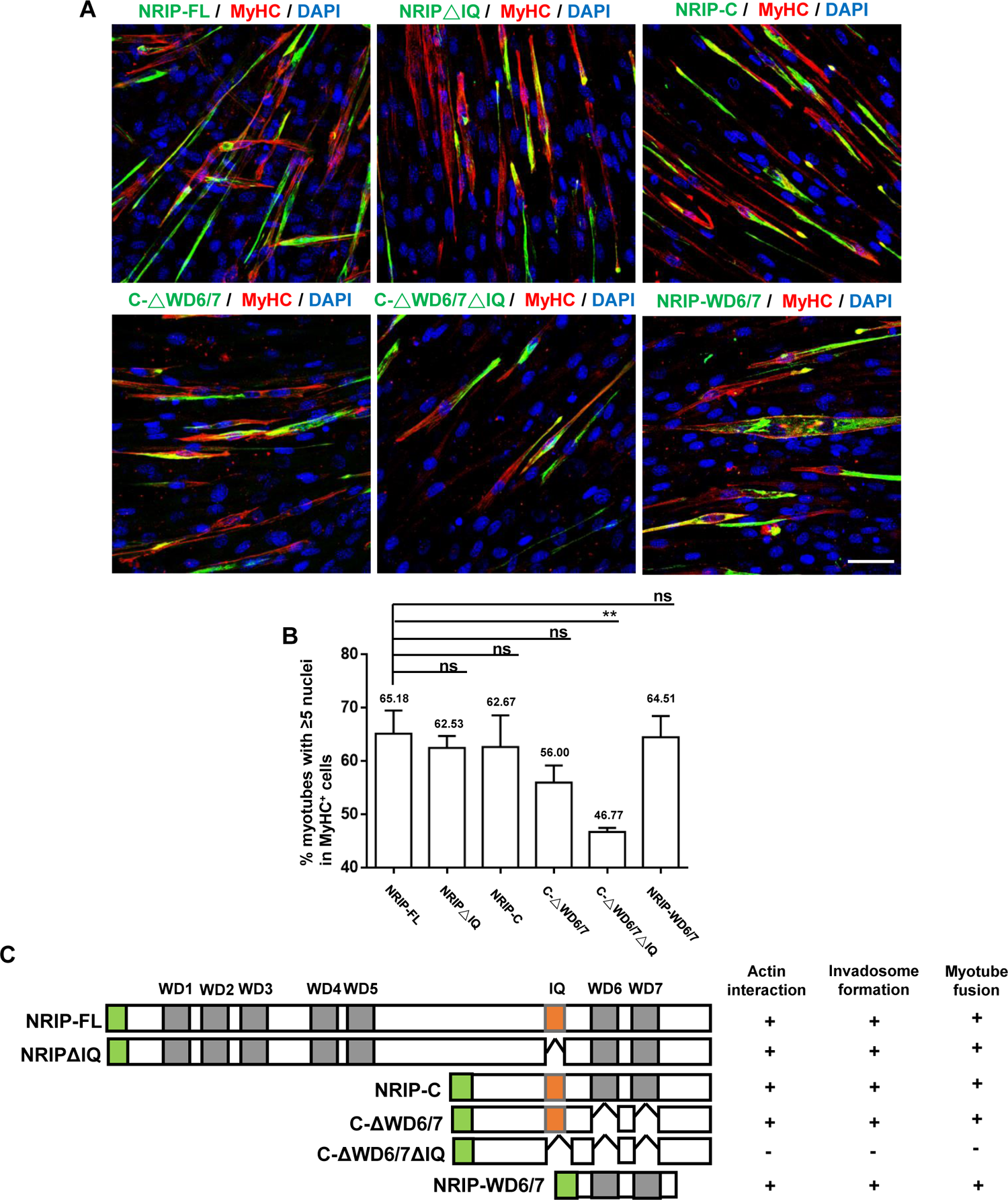
The correlation of NRIP-actin binding with myotube formation using NRIP mutant analysis. (A) The images of myotube formation. C2C12 myoblasts were transfected with NRIP mutants including NRIP-FL, NRIPΔIQ, NRIP-C, C-ΔWD6/7, C-ΔWD6/7ΔIQ and NRIP-WD6/7, then subjected to immunofluorescence stain with anti-EGFP (NRIP, green), and anti-MyHC antibody (red). DAPI for nuclear stain (blue). The C-ΔWD6/7ΔIQ loss of actin-binding reduced myotube size. Scale bar: 100 μm. (B) Quantitation of NRIP mutants for myotube formation. The percentage of myotubes (≥ 5 nuclei in MyHC^+^ cell) to total MyHC^+^ cells was measured from each NRIP mutant (N=6 for each group) and shown on top of the box. The C-ΔWD6/7ΔIQ showed reduced MyHC^+^ cell compared to NRIP-FL. (C) Summary of the relationship between actin-binding, invadosome formation and myotube fusion based on Figure 9 and 10. Data are mean ± SEM from 6 separate experiments. \*\**P* < 0.01; student *t* test. See also Figure 10— figure supplement 1, Figure 10—source data 1. Source data 1: Numerical data for Figure 10B.

## Discussion

In this paper, we demonstrated the NRIP expression profiles in the embryonic development of mice; NRIP was ubiquitously expressed in various tissues at the embryonic stage (Figure 1A and 1B). When specifically illustrating NRIP expression during muscle development, the muscle NRIP level was visible at the embryonic stage but robustly peaked at postnatal day P1 and P7, with detectable expression at 6 weeks (Figure 1C). In combination with our previous report, the NRIP level is the same at 16 and 6 weeks (Chen et al., 2018), which implies that muscle NRIP maintains a constant level in postnatal development at least until age 16 weeks. The functions of NRIP in muscle are caused by NRIP directly interacting with calmodulin (CaM) in the presence of calcium, which affects skeletal muscle functions such as muscle force, calcium storage, and mitochondrial activities (Chen et al., 2015; Chen et al., 2018). Additionally, NRIP can bind to α-actinin 2 (ACTN2) to regulate the length of the Z-band and maintain the integrity of the sarcomere structure in heart muscles (Yang et al., 2019). Moreover, NRIP also directly binds with acetylcholine receptor (AChR) to form AChR complexes for neuron muscular junction formation (Tsai et al., 2021). Deprivation of NRIP in muscle not only causes muscle cell pathology and abnormal neuron muscular junction, but also decreases α−motor neuron number (Chen et al., 2018); indicating that muscle NRIP can retrogradely regulate motor neuron growth in the spinal cord. Hence, the deprivation of muscle NRIP will cause motor neuron degeneration (Chen et al., 2018). Collectively, NRIP was more highly expressed postnatally than at the embryonic stage, which suggests that NRIP is important for muscle function postnatally. The deficiency of muscle NRIP leads to skeletal muscle dysfunction, cardiomyopathy, and motor neuron degeneration.

However, low NRIP expression at the embryonic stage can be the reason behind global NRIP-knockout mice being fertile and vital (Chen et al., 2015). Furthermore, NRIP is highly induced after muscle injury, and NRIP-knockout mice exhibit delayed muscle repair coupled with impaired myogenic capacity after muscle injury (Chen et al., 2015). Muscle myogenesis can occur in the embryonic stage and in adult regeneration (Abmayr & Pavlath, 2012; Hernandez-Hernandez et al., 2017). High expression of muscle NRIP at the postnatal stage may also indicate that muscle NRIP might initiate the repair process to promote muscle regeneration after stress (such as injury). In contrast, the myomaker, a muscle-specific membrane protein, functions to promote myoblast fusion during myogenesis. The expression profile is highly and only expressed in skeletal muscles during mouse embryogenesis, with no detectable expression after postnatal day 7; therefore, myomaker-knockout mice cannot survive until postnatal day 7 (Millay et al., 2013). Hence, the expression pattern of each protein can be related to its physiological homeostasis.

Here, we illustrated that NRIP was essential in myogenesis differentiation or myoblast fusion. As shown in Figures 2 and 3, the primary myoblasts from muscle-specific NRIP-knockout mice and NRIP-null C2C12 cells (KO19) showed significantly defective differentiation and fusion indexes, which suggests that NRIP is required for muscle myogenesis. Consistent with our previous report (Chen et al., 2015), NRIP-deficient mice were found to have exhibited a high proportion of small myofibers during muscle regeneration. In general, embryonic muscle formation and adult muscle regeneration include the activation and proliferation of satellite cells, myocyte differentiation, cell–cell fusion, and myotube formation (Abmayr & Pavlath, 2012; Hernandez-Hernandez et al., 2017). For myogenesis differentiation, myogenic regulatory factors, such as MyoD and myogenin, are transcriptional factors controlling the differentiation of skeletal muscle cells during embryogenesis and postnatal myogenesis (Hernandez-Hernandez et al., 2017). From the Pax7 and MyoD expression patterns, we found that NRIP functions in myogenesis differentiation; differentiated myocytes (Pax7^-^ MyoD^+^) of NRIP cKO mice appeared after 6 days as compared to WT mice, occurring 3 days after muscle injury (Figure 3-figure supplement 2). Thus, NRIP deficiency delays MyoD expression, thereby impairing differentiation and delaying myogenesis. Additionally, we previously reported that the mRNA and protein levels of myogenin are significantly lower in NRIP-KO mice than in WT mice at day 6 post-muscle injury (Chen et al., 2015), which implies that NRIP may regulate myogenin expression. Collectively, the loss of NRIP affects MyoD and myogenin expression. In contrast, myomaker (myoblast fusion gene) functions in myoblast fusion but not in myogenesis differentiation because MyoD and myogenin are expressed normally in myomaker embryos when comparing KO and WT mice (Millay et al., 2013). Taken together, NRIP deficiency delays the expression of MyoD and myogenin, resulting in impaired myogenesis differentiation. The mechanism by which NRIP regulates MyoD and myogenin remains a topic for future investigation.

NRIP plays a role in myoblast fusion (Figure 2 and 3). NRIP is a novel actin-binding protein (Figure 5) and was required for invadosome formation through actin binding (Figure 6 and 7). In terms of NRIP involved in myoblast fusion, both primary myoblasts from NRIP-KO mice and deficient C2C12 cells (KO19 cells) showed reduced myotube formation (Figure 2 and 3); overexpressed NRIP in KO19 cells could rescue myotube formation (Figure 4), and the forced expression of NRIP in C2C12 cells increased multinucleated myofiber size (Figure 4H, right panel and Figure 4-figure supplement 1B). All this evidence supports the idea that NRIP plays a role in myotube formation and increasing myofiber size.

NRIP is an actin-binding protein involved in invadosome formation for myotube formation, and actin is a major component of invadosome that plays a role in myoblast-myoblast or myoblast–myotube fusion. In addition to NRIP’s interaction with actin, we also demonstrated that NRIP colocalized with other invadosome components such as Tks5 (invadosome scaffold protein) and cortactin (actin binding protein; as an invadosome marker) at the tips of C2C12 myotubes (Figure 6B) (Chuang et al., 2019; Deng et al., 2015); indicating that NRIP may be involved in invadosome formation through actin binding to regulate myoblasts fusion. Several actin-binding proteins have been reported to play important roles in myoblast fusion, functioning in actin polymerization or actin crosslinking (Abmayr & Pavlath, 2012; Kim et al., 2015). An invadosome located at the tip of a fusion-competent myoblast contains enriched actin. Additionally, the interface between two fusion partners includes trans-interacting cell adhesion molecules. The invadosome from the fusion-competent myoblast is called a fusogenic synapse (Chen, 2011; Chuang et al., 2019; Kim et al., 2015; Sens et al., 2010). Genetic, cell biological, and electron microscopy studies have demonstrated that F-actin mediates cell fusion. Actin polymerization controls the fusion process between fusion-competent myoblasts and founder cells. During myoblast fusion, the asymmetrical distribution of the F-actin-enriched invadosome is important for invasion and fusion pore formation (Chuang et al., 2019; Sens et al., 2010). Here, we characterized the NRIP located at enlarged actin foci accumulated at the cell contact site of a myoblast and myotube (Figure 6E). Furthermore, this NRIP could form an invadosome during cell–cell fusion to proceed with myoblast fusion and myotube formation (Figs. 7 and 8), which indicates that NRIP is involved in invadosome formation through actin binding to trigger myoblast fusion.

To further support NRIP as a new component of an invadosome to affect cell–cell fusion via actin, time-lapse microscopy analysis revealed the dynamic changes of the actin cytoskeletal remodeling between NRIP and actin during myoblast fusion: at the start (0 min), the actin and NRIP-enriched foci remained in the cytoplasm; then, at 30–90 min, foci protruded toward the cell membrane to form a protrusive invadosome, which further confirms that NRIP participates in invasion during myoblast fusion (Figure 7 and Figure 7-figure supplement video 1). The asymmetry of F-actin in fusion-competent myoblasts that accumulates at the site of fusion is reportedly mediated by several components, such as actin-binding proteins Arp2/3, SCAR, dynamin-2 (Berger et al., 2008; Chuang et al., 2019; Kim et al., 2015) or the cell surface molecule M-cadherin, to form an M-cadherin/Rac1/Trio fusion complex for membrane curvature to regulate myoblast fusion (Abmayr & Pavlath, 2012; Chen & Olson, 2004; Kim et al., 2015). All these components can generate a dense F-actin focus to form an invadosome. Here, we showed that NRIP accumulated at the tips of differentiated myoblasts and formed an invadosome with actin (Figure 6) to form large myotubes containing multiple nuclei between myoblasts and nascent myotubes. Collectively, NRIP interacts with F-actin to form an invadosome focus at the fusion competent cells when cells start to fuse and then elongate over time during fusion, which further supports the notion that NRIP interacts with actin to form an invadosome to trigger myoblast fusion.

Additionally, NRIP consists of 860 amino acids, with seven WD40 repeats and one IQ motif (Chang et al., 2011; Tsai et al., 2005). To determine which domain of NRIP was responsible for actin binding and correlated with invadosome formation and myotube formation, NRIP mutation analysis revealed NRIP WD40 domains located at C fragment (named WD6/7) was responsible for actin binding, resulting in invadosome formation (Figure 9) and multinucleate myotube formation (Figure 10). WD40 domains are among the most abundant domains for protein–protein interaction; they form 7-blade-shaped structures and are exhibited as β-propeller architectures (Xu & Min, 2011). Proteins containing the WD40 domain can bind to several proteins to regulate various cellular functions, such as cytoskeletal assembly, signal transduction, vesicular trafficking, and transcription regulation (Xu & Min, 2011). Several WD40-containing proteins have been identified as actin-binding proteins; for example, coronin containing five WD40 domains interacts with actin to promote endocytosis and cell motility (Eckert et al., 2011); AIP1 (actin-interacting protein 1) containing two 7-blade WD40 β-propellers interacts with actin to promote the disassembly of actin filaments (Nomura et al., 2016). Here, NRIP containing seven WD40 domains (Tsai et al., 2005) could interact with actin through its C-terminal WD40 domain (WD6/7); WD6/7 only could significantly involve in invadosome formation (Figure 9) and myotube formation (Figure 10). On the other hand, NRIP is an α-actinin 2 (ACTN2) binding protein through its IQ domain (Yang et al., 2019) that can interact with actin for actin bundling formation (Winkelman et al., 2016). Here, the NRIP mutant, C-ΔWD6/7 (C fragment without WD40 domains but containing only one IQ motif), could indirectly interact with actin (Figure 9C and 9D) through ACTN2, which had the ability to form invadosomes (Figure 9E) and promote myotube formation (Figure 10), suggesting that the NRIP IQ domain could participate in myoblast fusion through ACTN2-actin binding. This further supports the idea that NRIP is a novel actin-binding protein for invadosome formation to induce myoblast fusion through its C-terminal WD40 domains or its IQ domain.

NRIP seems to be a multifunctional protein; its function might depend on its subcellular location. As shown in Figure 5, NRIP was identified as a membrane protein from its subcellular location by immunofluorescence analysis with phalloidin staining, and biochemical fractioning revealed approximately 50% of NRIP was present in the membrane fraction. This result supports our previous finding of NRIP colocalizing with acetylcholine receptors (AChRs) at the neuron muscular junction (Chen et al., 2018). In addition, NRIP is located at the cytosol (Figure 5D), which is consistent with our previous report of NRIP interacting with CaM or ACTN2 located at the cytosol sarcomere Z-disc to activate downstream calcineurin (CaN) and CaMKII signals or to stabilize Z-disc structure integrity, respectively (Chen et al., 2015; Yang et al., 2019). CaN-dependent signaling has been found to be involved in myoblast differentiation and skeletal muscle growth and hypertrophy (Friday et al., 2000; Park et al., 2016). Moreover, we found a nucleus location for NRIP (Figure 5E), which was previously characterized as a nucleus protein that interacts with the androgen receptor (AR) and serves as a transcriptional cofactor to drive AR-targeted gene expression (Chen et al., 2008; Tsai et al., 2005). Hence, the NRIP’s subcellular location may guide it to various biological and physical functions. Similarly, β-catenin, when in the nucleus as a transcriptional coactivator, forms a β-catenin/TCF/LEF-1 complex to control key developmental gene expression programs. In the absence of Wnt, cytoplasmic β-catenin protein is constantly degraded by the action of the Axin complex, including glycogen synthase kinase 3 (GSK3). However, the cadherin– catenin complex also plays a central role for β-catenin in this adhesion complex for cytoskeleton stability (Fu et al., 2008; MacDonald et al., 2009). Future studies could investigate the subcellular location of NRIP and associate its individual function with its subcellular location in different tissues and further investigate the mechanism of NRIP-regulating MyoD or myogenin via transcriptional activation in the nucleus or CaM-CaN signaling in the cytosol.

In sum, NRIP deficiency in mice or C2C12 cells affected both differentiation and fusion. Overexpressed NRIP in KO19 cells could rescue myotube formation, and the forced expression of NRIP in C2C12 cells increased myofiber size. Moreover, the loss of NRIP delayed MyoD expression, a marker of myogenesis differentiation, which supports NRIP involvement in differentiation. Intriguingly, NRIP was found to be a novel actin-binding protein and involved in the formation of a protrusive invadosome for an increased proportion of fused myotubes with more nuclei. Furthermore, NRIP was identified for direct participation in the fusion process by using a cell–cell fusion assay. NRIP mutation analysis demonstrated that NRIP interacted with actin either through WD40 domains for direct binding or indirectly through IQ domain for ACTN2 binding with actin, and the actin binding was correlated with between invadosome–myotube formation and actin-binding ability. Thus, NRIP affects myogenesis differentiation and acts as a novel actin-binding protein to form an invadosome for myotube fusion to form large-sized myofibers.

## Materials and Methods

### NRIP RNA *in situ* hybridization

NRIP RNA *in situ* hybridization was performed with two DIG-labeled antisense NRIP RNA probes (antisense probe 1-targeted exon 1 to 7, length 696 bp; antisense probe 2-targeted exon 14 to 19, length 578 bp) or one sense probe (sense probe-targeted exon 1 to 7, length 696 bp). The probe DNA process involved polymerase chain reaction (PCR) amplification with two pairs of primers (forward: 5’-CTACCTGGGAAGAAGAGAAT-3’, reverse: 5’-TGTAACACAGTGATGTCACT-3’ and forward: 5’-TGCAAGATACCGAACAGG-3’, reverse: 5’-CATCCTCATTTTCATTCTCC-3’) to produce two NRIP DNA fragments of 696 bp and 578 bp. Each NRIP DNA fragment was cloned into a pEASY-T3 vector (TransgeneBiotek, Telangana State, India) for NRIP RNA probe generation by *in vitro* transcription with SP6 RNA polymerase and T7 RNA polymerase, respectively, and labeled with digoxigenin (DIG) according to the DIG RNA Labeling Kit (SP6/T7). For RNA *in situ* hybridization, E11.5 mouse embryos were fixed with 4% paraformaldehyde (PFA) and hybridized with each probe, then detected via an alkaline phosphatase (AP)-conjugated anti-digoxigenin antibody, followed by a reaction of purple AP substrate for color development. The dark purple represents a positive signal showing the expression of the NRIP transcript.

### Primary myoblast isolation

The hindlimb muscles from 6-week-old NRIP cKO and WT mice were dissected and minced into small pieces. After digestion with enzymes (500 U/ml collagenase type II, 1.5 U/ml collagenase D, and 2.5 U/ml dispase II) for 60 min in a 37°C water bath and centrifugation at 300 × g for 5 min, the cell pellets were resuspended in the proliferation medium (DMEM containing 20% fetal bovine serum (FBS), 10% horse serum, 0.5% chicken embryo extract, and 2.5 ng/ml basic fibroblast growth factor [bFGF]), then seeded on dishes coated with Matrigel (0.09 mg/ml, CORNING) at 20% surface coverage and incubated at 37°C and 10% CO_2_ to allow for the attachment and migration of cells. Once local confluence was observed, cells were detached using 0.25% trypsin and seeded on dishes coated with collagen (0.1 mg/ml, CORNING). Cells were then cultured at 37°C and 10% CO_2_ for 1 h for attachment of fibroblasts and to leave myoblasts in the supernatant. The suspended myoblasts were then transferred to Matrigel-coated dishes for myoblast proliferation. For myoblast differentiation, cells were shifted to a differentiation medium (DMEM containing 5% horse serum) and cultured for 3 days to form multinucleated myotubes.

### Generation of NRIP-knockout C2C12 myoblasts

The CRISPR Cas9 target sites were identified at Zhang Lab’s website (http://crispr.mit.edu/), and the pair of all-in-one plasmid constructs (L-Cas9n-EGFP and R-Cas9n-puro) were obtained from Cold Spring Biotech Corp. The two guide RNAs (gRNAs) for NRIP genome targeting were expressed by L-Cas9n-EGFP and R-Cas9n-puro, and the sequences of single-guide RNAs (sgRNAs) were NRIP-sgRNA-F primer: 5’-GCC CGC ACC UGU UGU GGG AC and NRIP-sgRNA-R: 5’-CUU GGG CUG GAG GAC CCG UCC. The C2C12 cells were transfected with 2 μg plasmids (L-Cas9n-EGFP:R-Cas9n-puro=3:1) by electroporation (1650 v/10 ms/3 pulses) with the Neon Transfection System Kit (Thermo Fisher Scientific). The cells were seeded overnight to 24-well plates containing an antibiotic-free complete mediums. Cells were then selected with 3 μg/ml puromycin for 2 days and then harvested. Some cells were analyzed for editing efficiency, and others were diluted to 1 cell/well in 96-well plates for single cell culture for the period of 1 month. Half of the cells in each well were used to purify genomic DNA via the Epicentre MasterPure DNA Purification Kit (Epicentre). PCR was used to amplify the CRISPR reaction locus with designed primer pairs, and then each PCR fragment was cloned into the T3 cloning kit (ZGene Biotech) for DNA sequencing.

### Cell culture and transfection

C2C12, KO19 (NRIP-null C2C12 cells) and 293T cells were cultured in DMEM supplemented with 10% FBS. For C2C12 and KO19 differentiation, the culture medium was shifted to the differentiation medium, which consists of DMEM supplemented with 2% horse serum. For plasmid transfection, 293T cells were transfected with expression plasmids using jetPRIME (Polyplus), as recommended by the manufacturer. For C2C12 and KO19 transfection, cells were transfected with target plasmids using the K2 transfection kit (Biontex) following the manufacturer’s instructions.

### Immunofluorescence staining

Cells were fixed with 2% paraformaldehyde (PFA), which was followed by a permeabilization of −20 °C methanol. Cells were then blocked with 2% BSA for 30 min at room temperature. Samples were then incubated with the indicated primary antibodies [anti-NRIP (GeneTex, 1:200), anti-MyHC (Abcam, 1:200), anti-DsRed (TaKaRa, 1:200), anti-F-actin (Abcam, 1:200), anti-Tks5 (Santa Cruz, 1:100), anti-Cortactin (Santa Cruz, 1:100), and anti-GFP (Santa Cruz, 1:20)] at 4°C overnight. Then, the samples were incubated with fluorescent secondary antibodies (Cy3-conjugated mouse anti-rabbit, or 488-conjugated goat anti-mouse or 488-conjugated goat anti-rabbit, Jackson ImmunoResearch Laboratories) and mounted in DAPI Fluoromount-G (SouthernBiotech). The cell membrane was labeled with phalloidin (Sigma, 1:1000) for 1 h at room temperature. For tissue immunofluorescence analysis, tibialis anterior (TA) muscles were harvested and embedded in paraffin; then, tissue blocks were sectioned into 10-µm-thick slices and then incubated with boiled citrated acid (0.1M, pH 6.0) for antigen retrieval. Tissues were then blocked in 5% bovine serum albumin (BSA) and 1% horse serum for 1.5 h at room temperature and incubated with primary antibodies [anti-Pax7 (rabbit-polyclonal, Abcam, 1:500 dilution) and anti-MyoD (mouse-monoclonal, Millipore, 1:250 dilution)] overnight at 4°C. After the primary antibody reaction, sections were incubated with secondary antibodies. Finally, the sections were washed with PBS and mounted with DAPI Fluoromount-G (SouthernBiotech). Immunofluorescence signals were visualized and recorded using a Zeiss Axioskop 40 Optical Microscope with an AxioCam 702 camera and Zeiss Zen Blue software.

### Western blot analysis

Tissue and cell lysates were extracted and subjected to a western blot analysis. The primary antibodies were anti-NRIP (Novus, 1:2000), anti-EGF receptor (Cell Signaling, 1:1000), anti-DsRed (TaKaRa, 1:10000), anti-actin (Abcam, 1:10000), anti-GFP (Abcam, 1:10000), anti-flag (Abcam, 1:10000), anti-MyHC (Abcam, 1:2000), anti-Tks5 (Santa Cruz, 1:1000), anti-Cortactin (Santa Cruz, 1:1000) and anti-GAPDH (AbFrontier, 1:10000).

### His-MBP-NRIP purification and *in vitro* sedimentation assays

The His-MBP and His-MBP-NRIP proteins were expressed in the *E. coli* Rosetta strain (Novagen) by isopropyl β–D-1-thiogalactopyranoside (IPTG, 1 mM) induction. The bacterial lysates were harvested to collect the supernatants. The crude extracts were loaded into a column containing amylose resin (E8021S, New England Biolabs) for affinity chromatography. His-MBP and His-MBP-NRIP proteins were then eluted with binding buffer containing 10 mM maltose and concentrated in dialysis buffer (30 mM Tris, 500 mM NaCl) using AmiconR Ultra-15 Centrifugal Filter Devices. In the sedimentation assay, F-actin from bovine cardiac muscle (AD99, Cytoskeleton) and purified His-MBP-NRIP mixed in polymerization buffer (100 mM Tris pH 7.5, 500 mM KCl, 20 mM MgCl2, 10 mM ATP) were incubated for 1 h at room temperature. After centrifugation, the supernatant was removed, and pellets were resuspended in Milli-Q water and subjected to western blot analysis. In the His-pull down assay, His-MBP-NRIPs (NRIP-FL, NRIPΔIQ, NRIP-C, NRIP-C-ΔWD6/7 and NRIP-WD6/7) from bacteria were incubated with F-actin (AD99, Cytoskeleton) in the binding buffer (10 mM Tris-HCl, pH 8.0, 120 mM NaCl, 1 mM EDTA, 0.1% NP40, and protease inhibitor cocktail) at 4°C for 2 h. The protein mixtures were purified with Ni-NTA beads and subjected to western blot analysis.

### Immunoprecipitation assay

The 293T cells were co-transfected with pFlag-NRIP and pmCherry-actin, and then protein lysates were extracted and subjected to immunoprecipitation assay. For domain mapping of NRIP for actin binding, pmCherry-actin was co-transfected with pEGFP-NRIP-Full or its mutants (pEGFP-NRIP-C, pEGFP-NRIP△IQ, pEGFP-NRIP-C-△ WD6/7 and Flag-NRIP-WD6/7) (Chang et al., 2011; Tsai et al., 2005) into 293T cells, and the immunoprecipitation assay was performed and subjected to western blot analysis. For construction of pEGFP-NRIP-CΔWD6/7ΔIQ, pEGFP-NRIP-CΔWD6/7 was used as a template to amplify the target product with the indicated primers: forward 5’-GAATTGGATACTTTGAACATTAGACC, reverse 5’-ATCACCAGGTCCTGCTCGAT to delete IQ motif on pEGFP-NRIP-C-ΔWD6/7. For *in vitro* NRIP and actin binding, the purified His-MBP-NRIPs were incubated with F-actin (AD99, Cytoskeleton) in 1 ml of IP buffer (50 mM Tris-HCl, pH 8.0, 150 mM NaCl, 2 mM EDTA, 1% Triton X-100, and protease inhibitor cocktail) at 4°C for 1 h. The immunoprecipitation assay was performed with anti-F-actin antibody (Abcam, 2μg) and subjected to western blot analysis.

### Cytosolic and membrane protein fractionation

Cytosolic and membrane proteins were extracted using the Mem-Plus Membrane Protein Extraction Kit (Thermo). Briefly, cell lysates were harvested in growth medium and centrifuged at 300 × g for 5 min then washed with Cell Wash Solution, followed by lysing the lysates with a permeabilization buffer for 10 min with constant mixing at 4°C and centrifugation for 15 min at 300 × g to obtain the cytosolic fraction. Then, a solubilization buffer was added to cell pellets for 30 min through constant mixing and centrifugation at 16,000 × g for 15 min to extract solubilized membrane and membrane-associated proteins. The protein fractions of cytoplasm and membrane were analyzed by western blot assay.

### Cell fusion assay

C2C12 cells were transfected with pmCherry and KO19 transfected with pEGFP vector or pEGFP-NRIP by using the K2 Transfection kit (Biontex) for 24 h followed by replacement with fresh growth medium and culture for 24 h. Then, the cells were trypsinized and mixed in a ratio of 1:1 (each cell line: 7.1×10^4^ cells/ cm^2^) and seeded in 12-well plates covered by a cover glass. The next day, the cells were shifted to differentiation medium and incubated for 12 days to form fused myotubes; then, they were subjected to immunofluorescence staining.

### Time-lapse microscopy

C2C12 myoblasts were grown to 80% confluence and cotransfected with pmCherry-actin and pEGFP-NRIP by using a K2 Transfection kit (Biontex) for 24 h. They were then shifted to a differentiation medium for another 24 h. The cells were trypsinized and replated on Matrigel-coated cover glass to reach the proper cell density. Cells were replaced with imaging medium (phenol red-free differentiation medium), and a Laser TIRF/Spinning Disc Confocal Microscope was used for time-lapse imaging at 37°C, 5% CO_2_ with time intervals of 10 min for 6 h.

### Statistical analysis

All statistical data were analyzed using Prism (GraphPad Software). Data are presented as mean ± SEM. Two groups were compared by the student’s t test. *P* < 0.05 was considered statistically significant.

### Study approval

All animal procedures were reviewed and approved by the Institutional Animal Care and Usage Committee (IACUC) of the College of Medicine, National Taiwan University. The experimental mice were housed in the animal center under a 12-h light/dark cycle with free access to food and water.

### Online supplemental materials

Figure supplement 1 shows the scheme of generating NRIP-null cell lines by the CRISPR-Cas9 system. Figure supplement 2 presents loss of NRIP reduces satellite cell activation and delayed regeneration in injured muscles. Figure supplement 3 shows the percentage of myonuclei number distribution of MyHC^+^ cells in NRIP-overexpressed KO19 and C2C12 cells. Figure supplement 4 shows the comparable expression level of MyHC protein among C2C12 myotubes transfected with NRIP mutants. Figure supplement video 1 shows the time-lapse microscopy for NRIP and actin during the formation of a podosome-like structure (Figs. 10B and 10C).

## Acknowledgements

We appreciate Scribendi Inc. for their English editing. Additionally, muscle-specific NRIP knockout mice (cKO) were bred in the Transgenic Mouse Core Facility at the National Taiwan University Center for Genomic Medicine. We thank the staff of the Gene Knockout/in Cell Line Modeling Core at the First Core Labs, National Taiwan University College of Medicine, for generating NRIP-null C2C12 myoblasts (KO19); the confocal image analysis at The Imaging Core at the First Core Lab, National Taiwan University College of Medicine, and the Microscopy Core Facility, Department of Medical Research, National Taiwan University Hospital; and the paraffin preparation section of the Animal Center, National Taiwan University College of Medicine. We thank Drs. Shiou-Ru Tzeng and Ya-Wen Liu for the paper discussion, and the plasmid-Lifeact-RFP was provided by Dr. Ya-Wen Liu. This work was supported by the Ministry of Science and Technology [MOST 104-2320-B-002-053-MY3 to S-L. Chen, MOST 107-2320-B-002-012-MY3 to S-L. Chen, MOST 110-2320-B-002-052-MY3 to S-L. Chen] and the National Health Research Institute [NHRI-105-1053ISI to S-L. Chen].

The authors declare no competing financial interests.

## Authors’ contributions

S-L. Chen, H-H. Chen and Y-J. Han conceived and designed the experiments. H-H. Chen., Y-J. Han, T-C. Wu, W-S. Yen, T-Y. Lai, and P-H. Wei performed the experiments. H-H. Chen, Y-J. Han, T-C. Wu, W-S. Yen, T-Y. Lai, P-H. Wei, L-K. Tsai, H-J. Lai, Y-P. Tsao and S-L. Chen analyzed the data. S-L. Chen contributed the reagents, materials, and analysis tools. S-L. Chen, H-H. Chen, Y-J. Han, T-C. Wu, and W-S. Yen contributed to writing the manuscript.

## SUPPLEMENTARY FIGURE LEGENDS

**Figure 3 - Figure Supplement 1.**
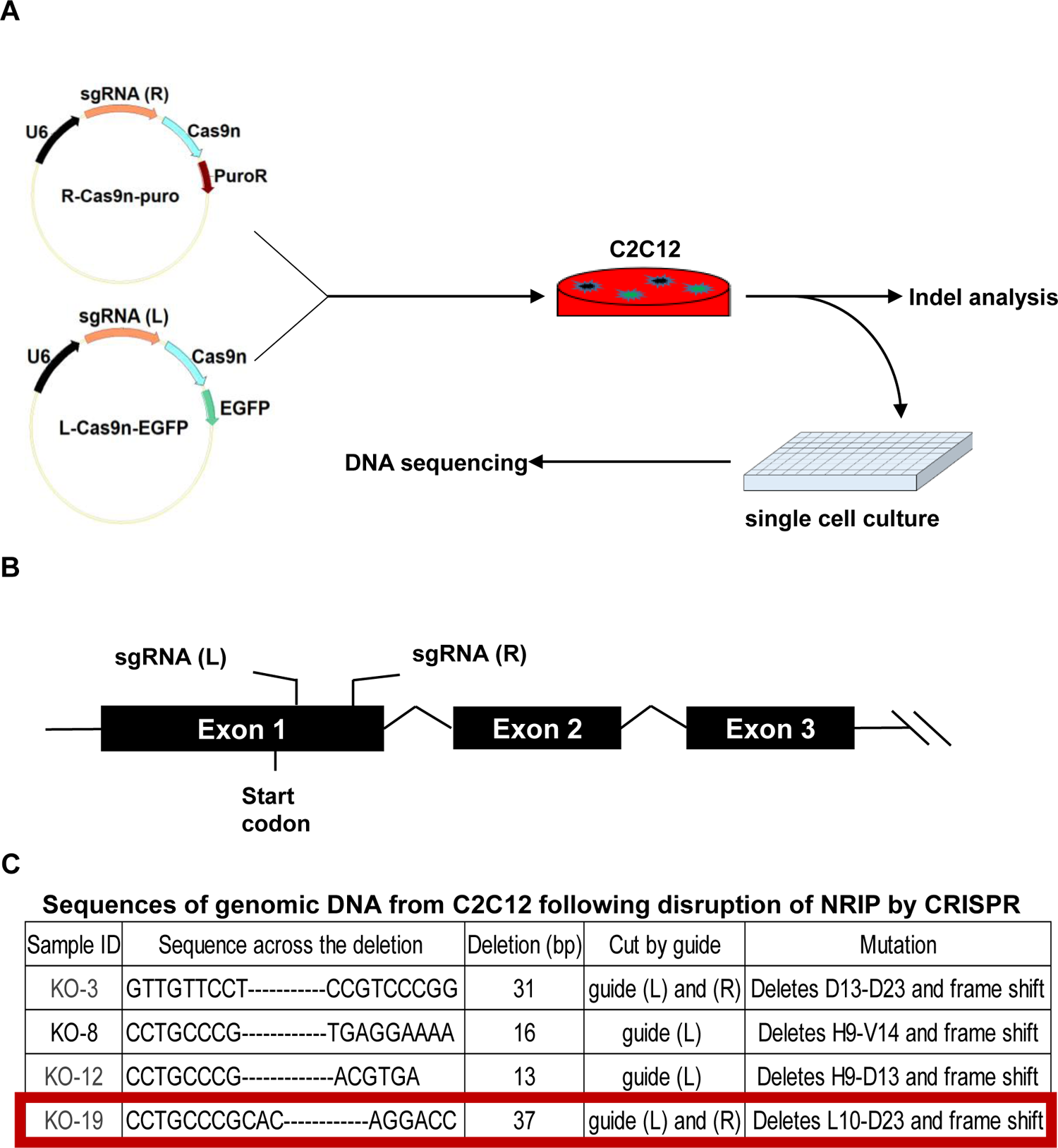
Generation of NRIP-null C2C12 cells. (A) Schematic representation of the generation of NRIP-null cell lines via the CRISPR-Cas9 system. Trypsinized C2C12 cells (1×10^5^ cells) were added with 2 µg plasmids (sgRNA-L-Cas9n-EGFP: sgRNA-R-Cas9n-puro = 3:1) and subjected to electroporation (1650 v/10 ms/3 pulses). Cells were seeded into 24-well plates containing antibiotic-free complete mediums, which were incubated overnight, and then selected with 3 µg/ml puromycin for 2 days. Then, the cells were diluted to 1 cell per well for single-cell cultures. After 1 month, genomic DNA was harvested from half the cells per well for DNA sequencing. (B) The location of sgRNAs for NRIP gene editing. NRIP-sgRNA-F primer: GCC CGC ACC UGU UGU GGG AC; NRIP-sgRNA-R: CUU GGG CUG GAG GAC CCG UCC. The NRIP gene consists of 19 exons with the translation starting site at exon 1. NRIP exon 1 was targeted by pairs of sgRNAs. (C) Sequences of genomic DNA from NRIP-null C2C12 cell lines and the locations of deletion sequences at the targeted NRIP from four NRIP-null cell lines. KO19 cells were chosen for the following assay.

**Figure 3 - Figure Supplement 2.**
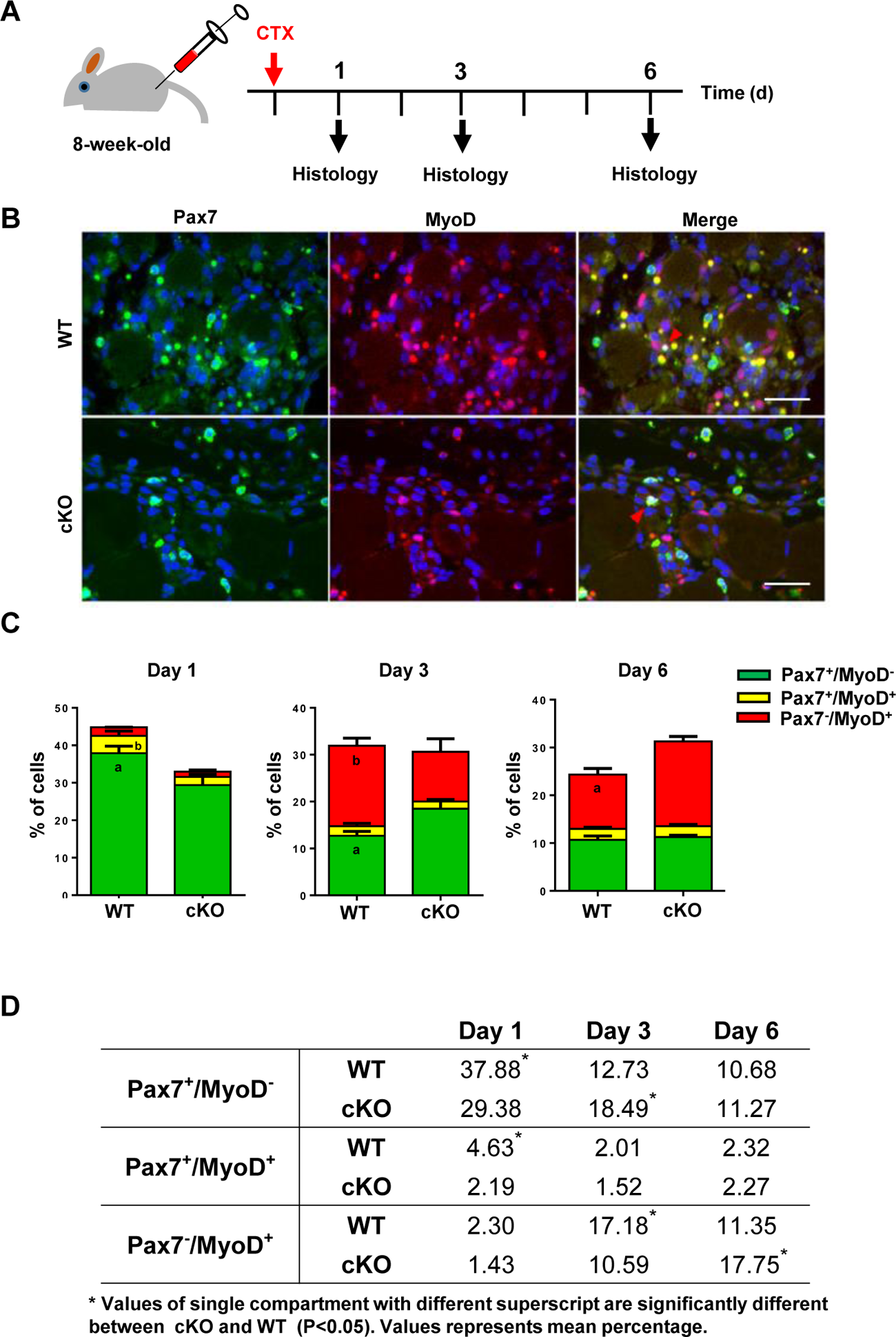
The deficiency of muscle NRIP delays myoblast activation and differentiation during muscle regeneration. (A) Schematic representation of cardiotoxin (CTX) treatment in 8-week-old male WT mice and cKO mice. The mice’s tibialis anterior (TA) muscles were injected with a single dose of 10 mM cardiotoxin and harvested for histological analysis at 1, 3, and 6 days after treatment. (B) Representative immunofluorescence images of the TA muscles at 3 days after injury. Paraffin sections were stained with anti-Pax7 (green), anti-MyoD (red) antibodies, and DAPI (blue, nucleus stain). Scale bars: 100 µm. (C) Quantification of Pax7^+^/MyoD^-^ (green box), Pax7^+^/MyoD^+^ (yellow box), and Pax7^-^/MyoD^+^ cells (red box) at days 1, 3, and 6 after injury in the TA muscles. N = 6 per genotype, 600 cells counted for each TA muscle. (D) The percentage distribution of Pax7^+^/MyoD^-^, Pax7^+^/MyoD^+^, and Pax7^-^/MyoD^+^ cells at 1, 3, and 6 days after injury in the tibialis anterior muscles of WT and NRIP cKO mice. N = 6 per genotype; 600 cells counted for each TA muscle. Data are mean ± SEM from 3 separate experiments. \**P* < 0.05

**Figure 4 - Figure Supplement 1.**
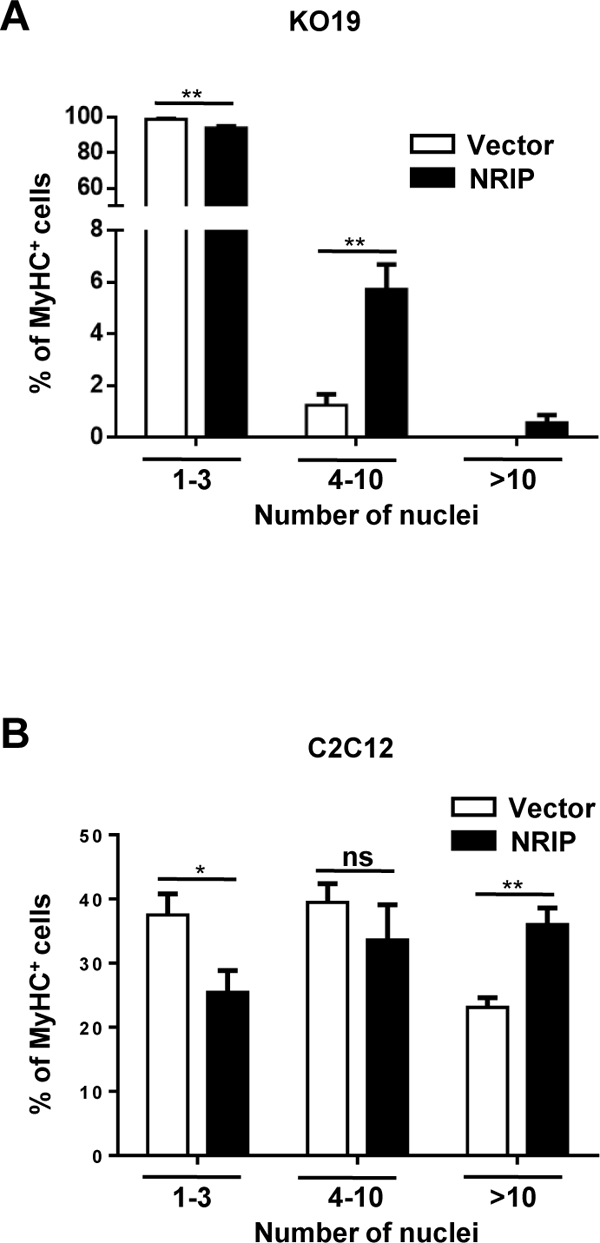
The percentage of myonuclei number distribution of MyHC^+^ cells. (A) NRIP in KO19 cells (Figure 4). (B) NRIP in C2C12 cells (Figure 4). Myonuclei numbers are categorized into three groups: 1–3 nuclei, 4–10 nuclei, and > 10 nuclei. Data are mean ± SEM from 3 separate experiments. \**P* < 0.05, \*\**P* < 0.01, ns, not significant as per the student’s t-test.

**Figure 10 - Figure Supplement 1.**
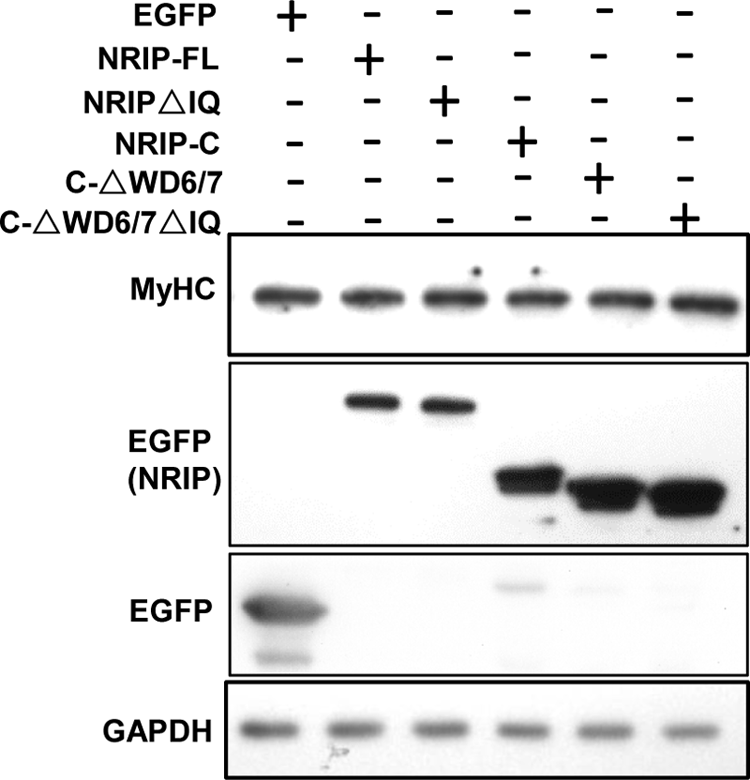
The comparable expression level of MyHC protein among C2C12 myotubes transfected with NRIP mutants. C2C12 myoblasts were transiently transfected with each EGFP-tagged NRIP mutant plasmid and differentiated for 3 days. The cell lysates were subjected to immunoblotting with anti-MyHC for MyHC protein expression, anti-EGFP for EGFP-tagged NRIP mutant expression, and anti-GAPDH for protein loading control. See also Figure 10—figure supplement 1-source data1. Source data 1: Source WB blot for Figure 10—figure supplement 1.

---

Figure 7 - Figure Supplement Video 1: Time-lapse microscopy for NRIP and actin during invadosome structure formation (Figure 7B).

## References

1. Abmayr, S. M., & Pavlath, G. K. (2012). Myoblast fusion: lessons from flies and mice. Development, 139(4), 641–656. https://doi.org/10.1242/dev.068353

2. Aline, G., & Sotiropoulos, A. (2012). Srf: A key factor controlling skeletal muscle hypertrophy by enhancing the recruitment of muscle stem cells. Bioarchitecture, 2(3), 88–90. https://doi.org/10.4161/bioa.20699

3. Bentzinger, C. F., Wang, Y. X., & Rudnicki, M. A. (2012). Building muscle: molecular regulation of myogenesis. Cold Spring Harb Perspect Biol, 4(2). https://doi.org/10.1101/cshperspect.a008342

4. Berger, S., Schafer, G., Kesper, D. A., Holz, A., Eriksson, T., Palmer, R. H., Beck, L., Klambt, C., Renkawitz-Pohl, R., & Onel, S. F. (2008). WASP and SCAR have distinct roles in activating the Arp2/3 complex during myoblast fusion. Journal of Cell Science, 121(8), 1303–1313. https://doi.org/10.1242/jcs.022269

5. Berkes, C. A., & Tapscott, S. J. (2005). MyoD and the transcriptional control of myogenesis. Semin Cell Dev Biol, 16(4-5), 585–595. https://doi.org/10.1016/j.semcdb.2005.07.006

6. Bi, P., Ramirez-Martinez, A., Li, H., Cannavino, J., McAnally, J. R., Shelton, J. M., Sanchez-Ortiz, E., Bassel-Duby, R., & Olson, E. N. (2017). Control of muscle formation by the fusogenic micropeptide myomixer. Science, 356(6335), 323–327. https://doi.org/10.1126/science.aam9361

7. Chang, S. W., Tsao, Y. P., Lin, C. Y., & Chen, S. L. (2011). NRIP, a novel calmodulin binding protein, activates calcineurin to dephosphorylate human papillomavirus E2 protein. J Virol, 85(13), 6750–6763. https://doi.org/10.1128/JVI.02453-10

8. Chen, E. H. (2011). Invasive podosomes and myoblast fusion. Curr Top Membr, 68, 235–258. https://doi.org/10.1016/B978-0-12-385891-7.00010-6

9. Chen, E. H., & Olson, E. N. (2004). Towards a molecular pathway for myoblast fusion in Drosophila. Trends Cell Biol, 14(8), 452–460. https://doi.org/10.1016/j.tcb.2004.07.008

10. Chen, H. H., Chen, W. P., Yan, W. L., Huang, Y. C., Chang, S. W., Fu, W. M., Su, M. J., Yu, I. S., Tsai, T. C., Yan, Y. T., Tsao, Y. P., & Chen, S. L. (2015). NRIP is newly identified as a Z-disc protein, activating calmodulin signaling for skeletal muscle contraction and regeneration. J Cell Sci, 128(22), 4196–4209. https://doi.org/10.1242/jcs.174441

11. Chen, H. H., Fan, P., Chang, S. W., Tsao, Y. P., Huang, H. P., & Chen, S. L. (2017). NRIP/DCAF6 stabilizes the androgen receptor protein by displacing DDB2 from the CUL4A-DDB1 E3 ligase complex in prostate cancer. Oncotarget, 8(13), 21501–21515. https://doi.org/10.18632/oncotarget.15308

12. Chen, H. H., Tsai, L. K., Liao, K. Y., Wu, T. C., Huang, Y. H., Huang, Y. C., Chang, S. W., Wang, P. Y., Tsao, Y. P., & Chen, S. L. (2018). Muscle-restricted nuclear receptor interaction protein knockout causes motor neuron degeneration through down-regulation of myogenin at the neuromuscular junction. J Cachexia Sarcopenia Muscle, 9(4), 771–785. https://doi.org/10.1002/jcsm.12299

13. Chen, J. C., & Goldhamer, D. J. (2003). Skeletal muscle stem cells. Reprod Biol Endocrinol, 1, 101. https://doi.org/10.1186/1477-7827-1-101

14. Chen, P. H., Tsao, Y. P., Wang, C. C., & Chen, S. L. (2008). Nuclear receptor interaction protein, a coactivator of androgen receptors (AR), is regulated by AR and Sp1 to feed forward and activate its own gene expression through AR protein stability. Nucleic Acids Res, 36(1), 51–66. https://doi.org/10.1093/nar/gkm942

15. Chuang, M. C., Lin, S. S., Ohniwa, R. L., Lee, G. H., Su, Y. A., Chang, Y. C., Tang, M. J., & Liu, Y. W. (2019). Tks5 and Dynamin-2 enhance actin bundle rigidity in invadosomes to promote myoblast fusion. J Cell Biol, 218(5), 1670–1685. https://doi.org/10.1083/jcb.201809161

16. Dalkilic, I., Schienda, J., Thompson, T. G., & Kunkel, L. M. (2006). Loss of FilaminC (FLNc) results in severe defects in myogenesis and myotube structure. Mol Cell Biol, 26(17), 6522–6534. https://doi.org/10.1128/MCB.00243-06

17. De Bari, C., Dell’Accio, F., Vandenabeele, F., Vermeesch, J. R., Raymackers, J. M., & Luyten, F. P. (2003). Skeletal muscle repair by adult human mesenchymal stem cells from synovial membrane. J Cell Biol, 160(6), 909–918. https://doi.org/10.1083/jcb.200212064

18. Deng, S., Bothe, I., & Baylies, M. K. (2015). The Formin Diaphanous Regulates Myoblast Fusion through Actin Polymerization and Arp2/3 Regulation. PLoS Genet, 11(8), e1005381. https://doi.org/10.1371/journal.pgen.1005381

19. Eckert, C., Hammesfahr, B., & Kollmar, M. (2011). A holistic phylogeny of the coronin gene family reveals an ancient origin of the tandem-coronin, defines a new subfamily, and predicts protein function. BMC Evol Biol, 11, 268. https://doi.org/10.1186/1471-2148-11-268

20. Friday, B. B., Horsley, V., & Pavlath, G. K. (2000). Calcineurin activity is required for the initiation of skeletal muscle differentiation. J Cell Biol, 149(3), 657–666. https://doi.org/10.1083/jcb.149.3.657

21. Fu, A. K., Cheung, Z. H., & Ip, N. Y. (2008). Beta-catenin in reverse action. Nat Neurosci, 11(3), 244–246. https://doi.org/10.1038/nn0308-244

22. Hernandez-Hernandez, J. M., Garcia-Gonzalez, E. G., Brun, C. E., & Rudnicki, M. A. (2017). The myogenic regulatory factors, determinants of muscle development, cell identity and regeneration. Semin Cell Dev Biol, 72, 10–18. https://doi.org/10.1016/j.semcdb.2017.11.010

23. Ishibashi, J., Perry, R. L., Asakura, A., & Rudnicki, M. A. (2005). MyoD induces myogenic differentiation through cooperation of its NH2- and COOH-terminal regions. J Cell Biol, 171(3), 471–482. https://doi.org/10.1083/jcb.200502101

24. Jin, J., Arias, E. E., Chen, J., Harper, J. W., & Walter, J. C. (2006). A family of diverse Cul4-Ddb1-interacting proteins includes Cdt2, which is required for S phase destruction of the replication factor Cdt1. Mol Cell, 23(5), 709–721. https://doi.org/10.1016/j.molcel.2006.08.010

25. Kang, J. S., & Krauss, R. S. (2010). Muscle stem cells in developmental and regenerative myogenesis. Curr Opin Clin Nutr Metab Care, 13(3), 243–248. https://doi.org/10.1097/MCO.0b013e328336ea98

26. Kim, J. H., Jin, P., Duan, R., & Chen, E. H. (2015). Mechanisms of myoblast fusion during muscle development. Curr Opin Genet Dev, 32, 162–170. https://doi.org/10.1016/j.gde.2015.03.006

27. Kim, S., Shilagardi, K., Zhang, S., Hong, S. N., Sens, K. L., Bo, J., Gonzalez, G. A., & Chen, E. H. (2007). A critical function for the actin cytoskeleton in targeted exocytosis of prefusion vesicles during myoblast fusion. Dev Cell, 12(4), 571–586. https://doi.org/10.1016/j.devcel.2007.02.019

28. Knight, J. D., & Kothary, R. (2011). The myogenic kinome: protein kinases critical to mammalian skeletal myogenesis. Skelet Muscle, 1, 29. https://doi.org/10.1186/2044-5040-1-29

29. MacDonald, B. T., Tamai, K., & He, X. (2009). Wnt/beta-catenin signaling: components, mechanisms, and diseases. Dev Cell, 17(1), 9–26. https://doi.org/10.1016/j.devcel.2009.06.016

30. Mattila, P. K., & Lappalainen, P. (2008). Filopodia: molecular architecture and cellular functions. Nat Rev Mol Cell Biol, 9(6), 446–454. https://doi.org/10.1038/nrm2406

31. Millay, D. P., O’Rourke, J. R., Sutherland, L. B., Bezprozvannaya, S., Shelton, J. M., Bassel-Duby, R., & Olson, E. N. (2013). Myomaker is a membrane activator of myoblast fusion and muscle formation. Nature, 499(7458), 301–305. https://doi.org/10.1038/nature12343

32. Miller, K. J., Thaloor, D., Matteson, S., & Pavlath, G. K. (2000). Hepatocyte growth factor affects satellite cell activation and differentiation in regenerating skeletal muscle. Am J Physiol Cell Physiol, 278(1), C174–181. http://www.ncbi.nlm.nih.gov/pubmed/10644525

33. Mimura, N., & Asano, A. (1986). Isolation and characterization of a conserved actin-binding domain from rat hepatic actinogelin, rat skeletal muscle, and chicken gizzard alpha-actinins. J Biol Chem, 261(23), 10680–10687. https://www.ncbi.nlm.nih.gov/pubmed/3733725

34. Nomura, K., Hayakawa, K., Tatsumi, H., & Ono, S. (2016). Actin-interacting Protein 1 Promotes Disassembly of Actin-depolymerizing Factor/Cofilin-bound Actin Filaments in a pH-dependent Manner. J Biol Chem, 291(10), 5146–5156. https://doi.org/10.1074/jbc.M115.713495

35. Nowak, S. J., Nahirney, P. C., Hadjantonakis, A. K., & Baylies, M. K. (2009). Nap1-mediated actin remodeling is essential for mammalian myoblast fusion. J Cell Sci, 122(Pt 18), 3282–3293. https://doi.org/10.1242/jcs.047597

36. O’Connell, M. E., Sridharan, D., Driscoll, T., Krishnamurthy, I., Perry, W. G., & Applewhite, D. A. (2019). The Drosophila protein, Nausicaa, regulates lamellipodial actin dynamics in a Cortactin-dependent manner. Biol Open, 8(6). https://doi.org/10.1242/bio.038232

37. Olguin, H. C., Yang, Z., Tapscott, S. J., & Olwin, B. B. (2007). Reciprocal inhibition between Pax7 and muscle regulatory factors modulates myogenic cell fate determination. J Cell Biol, 177(5), 769–779. https://doi.org/10.1083/jcb.200608122

38. Park, S. Y., Yun, Y., Lim, J. S., Kim, M. J., Kim, S. Y., Kim, J. E., & Kim, I. S. (2016). Stabilin-2 modulates the efficiency of myoblast fusion during myogenic differentiation and muscle regeneration. Nat Commun, 7, 10871. https://doi.org/10.1038/ncomms10871

39. Quinn, M. E., Goh, Q., Kurosaka, M., Gamage, D. G., Petrany, M. J., Prasad, V., & Millay, D. P. (2017). Myomerger induces fusion of non-fusogenic cells and is required for skeletal muscle development. Nat Commun, 8, 15665. https://doi.org/10.1038/ncomms15665

40. Richardson, B. E., Beckett, K., Nowak, S. J., & Baylies, M. K. (2007). SCAR/WAVE and Arp2/3 are crucial for cytoskeletal remodeling at the site of myoblast fusion. Development, 134(24), 4357–4367. https://doi.org/10.1242/dev.010678

41. Riedl, J., Crevenna, A. H., Kessenbrock, K., Yu, J. H., Neukirchen, D., Bista, M., Bradke, F., Jenne, D., Holak, T. A., Werb, Z., Sixt, M., & Wedlich-Soldner, R. (2008). Lifeact: a versatile marker to visualize F-actin. Nat Methods, 5(7), 605–607. https://doi.org/10.1038/nmeth.1220

42. Sens, K. L., Zhang, S., Jin, P., Duan, R., Zhang, G., Luo, F., Parachini, L., & Chen, E. H. (2010). An invasive podosome-like structure promotes fusion pore formation during myoblast fusion. J Cell Biol, 191(5), 1013–1027. https://doi.org/10.1083/jcb.201006006

43. Shahini, A., Vydiam, K., Choudhury, D., Rajabian, N., Nguyen, T., Lei, P., & Andreadis, S. T. (2018). Efficient and high yield isolation of myoblasts from skeletal muscle. Stem Cell Res, 30, 122–129. https://doi.org/10.1016/j.scr.2018.05.017

44. Tsai, L. K., Chen, I. H., Chao, C. C., Hsueh, H. W., Chen, H. H., Huang, Y. H., Weng, R. W., Lai, T. Y., Tsai, Y. C., Tsao, Y. P., & Chen, S. L. (2021). Autoantibody of NRIP, a novel AChR-interacting protein, plays a detrimental role in myasthenia gravis. J Cachexia Sarcopenia Muscle, 12(3), 665–676. https://doi.org/10.1002/jcsm.12697

45. Tsai, T. C., Lee, Y. L., Hsiao, W. C., Tsao, Y. P., & Chen, S. L. (2005). NRIP, a novel nuclear receptor interaction protein, enhances the transcriptional activity of nuclear receptors. J Biol Chem, 280(20), 20000–20009. https://doi.org/10.1074/jbc.M412169200

46. Winkelman, J. D., Suarez, C., Hocky, G. M., Harker, A. J., Morganthaler, A. N., Christensen, J. R., Voth, G. A., Bartles, J. R., & Kovar, D. R. (2016). Fascin- and alpha-Actinin-Bundled Networks Contain Intrinsic Structural Features that Drive Protein Sorting. Curr Biol, 26(20), 2697–2706. https://doi.org/10.1016/j.cub.2016.07.080

47. Xu, C., & Min, J. (2011). Structure and function of WD40 domain proteins. Protein Cell, 2(3), 202–214. https://doi.org/10.1007/s13238-011-1018-1

48. Yablonka-Reuveni, Z., Seger, R., & Rivera, A. J. (1999). Fibroblast growth factor promotes recruitment of skeletal muscle satellite cells in young and old rats. J Histochem Cytochem, 47(1), 23–42. http://www.ncbi.nlm.nih.gov/pubmed/9857210

49. Yang, K. C., Chuang, K. W., Yen, W. S., Lin, S. Y., Chen, H. H., Chang, S. W., Lin, Y. S., Wu, W. L., Tsao, Y. P., Chen, W. P., & Chen, S. L. (2019). Deficiency of nuclear receptor interaction protein leads to cardiomyopathy by disrupting sarcomere structure and mitochondrial respiration. J Mol Cell Cardiol, 137, 9–24. https://doi.org/10.1016/j.yjmcc.2019.09.009

50. Yin, H., Price, F., & Rudnicki, M. A. (2013). Satellite cells and the muscle stem cell niche. Physiol Rev, 93(1), 23–67. https://doi.org/10.1152/physrev.00043.2011

51. Zammit, P. S., Partridge, T. A., & Yablonka-Reuveni, Z. (2006). The skeletal muscle satellite cell: the stem cell that came in from the cold. J Histochem Cytochem, 54(11), 1177–1191. https://doi.org/10.1369/jhc.6R6995.2006

52. Zammit, P. S., Relaix, F., Nagata, Y., Ruiz, A. P., Collins, C. A., Partridge, T. A., & Beauchamp, J. R. (2006). Pax7 and myogenic progression in skeletal muscle satellite cells. J Cell Sci, 119(Pt 9), 1824–1832. https://doi.org/10.1242/jcs.02908

53. Zhang, Q., Vashisht, A. A., O’Rourke, J., Corbel, S. Y., Moran, R., Romero, A., Miraglia, L., Zhang, J., Durrant, E., Schmedt, C., Sampath, S. C., & Sampath, S. C. (2017). The microprotein Minion controls cell fusion and muscle formation. Nat Commun, 8, 15664. https://doi.org/10.1038/ncomms15664

